# Homeostasis of mRNA concentrations through coupling transcription, exportation, and degradation

**DOI:** 10.1101/2023.08.23.554439

**Authors:** Qirun Wang, Jie Lin

## Abstract

Many experiments showed that eukaryotic cells maintain a constant mRNA concentration upon various perturbations by actively regulating mRNA production and degradation rates, known as mRNA buffering. However, the underlying mechanism is still unknown. Here, we propose a mechanistic model of mRNA buffering: the releasing-shuttling (RS) model. The model incorporates two crucial factors, X and Y, which play key roles in the transcription, exportation, and degradation processes. The model explains the constant mRNA concentration under genome-wide genetic perturbations and cell volume changes. Moreover, it quantitatively explains the slowed-down mRNA degradation after Pol II depletion and the temporal transcription dynamics after Xrn1 depletion. The RS model suggests that X and Y are likely composed of multiple molecules possessing redundant functions. We also present a list of X and Y candidates, and an experimental method to identify X. Our work uncovers a possible coupling mechanism between transcription, exportation, and degradation.

## INTRODUCTION

Maintaining an appropriate mRNA level is vital for all living cells. Surprisingly, many eukaryotic cells adjust the mRNA production rate and the mRNA degradation rate to achieve a constant mRNA concentration, an elegant but mysterious phenomenon called mRNA buffering [1–3]. Multiple experiments carried out in the budding yeast *Saccharomyces cerevisiae* showed that the total mRNA concentration and also the mRNA concentrations of most individual genes are invariant against various genetic perturbations [4–13]. In cases where transcription-related genes are perturbed, the mRNA production rate decreases; interestingly, the mRNA degradation rate decreases accordingly so that the mRNA concentration is invariant. In cases where mRNA degradation-related genes are perturbed, the mRNA degradation rate drops, and the production rate decreases accordingly to maintain a constant mRNA concentration. Beyond the steady-state results, Chappleboim et al. recently showed that after an acute depletion of mRNA degradation factors, the mRNA degradation rate dropped immediately. However, the production rate decreased after a delay, so the total mRNA concentration temporarily accumulated and then gradually returned to its original level [14]. mRNA buffering phenomenon has been reported in mammalian cells as well [15, 16], suggesting its universality across eukaryotes. Strikingly, Berry et al. subjected mammalian cells to genome-wide genetic perturbation screening and found that the total mRNA concentration in both the nucleus and cytoplasm remained virtually unchanged despite a significant variation in the mRNA production rate [17].

Meanwhile, the total mRNA concentration and the mRNA concentrations of most genes are also constant as the cell volume increases [17–20]. The homeostasis of mRNA concentration in a growing cell volume is often considered a consequence of an mRNA production rate proportional to cell volume with a constant mRNA degradation rate [21, 22]. The volume-scaling mRNA production rate has been proposed to come from the limiting nature of Pol II [23, 24] – the mRNA production rate is proportional to the copy number of RNA polymerase II (Pol II), which is proportional to the cell volume. However, Mena et al. found that the mRNA production rate is constant as the cell volume increases in *Saccharomyces cerevisiae*; nevertheless, homeostasis of mRNA concentration is still maintained due to the decreased mRNA degradation rate [25]. Similarly, Swaffer et al. found that the mRNA production rate increases sublinearly with the cell volume in *Saccharomyces cerevisiae*. In the meantime, the mRNA degradation rate decreases accordingly and perfectly balances the mRNA production rate so that the mRNA concentration remains constant [26].

In all the above examples, a coupling between mRNA production and mRNA degradation exists and balances the two rates so that the mRNA concentration is constant. It has been suggested that mRNA feedback to its own production or degradation may lead to mRNA buffering [14, 17, 26]. In addition to mRNA itself, several proteins have been suggested as possible players in mRNA buffering. One notable protein is Xrn1, the 5’3’ exonuclease that plays a crucial role in cytoplasmic mRNA degradation. Recent studies have shown that it also shuttles into the nucleus and functions in transcription regulation, suggesting its potential role in connecting mRNA production and degradation [8, 9, 27–29]. Similarly, the Ccr4-Not complex involved in mRNA deadenylation has also been found to shuttle between cytoplasm and nucleus and regulate transcription [30]. Other examples include Snf1, the yeast ortholog of AMPactivated protein kinase (AMPK), which interacts with both degradation machinery and transcription factors [31], and Rpb4/7, which are subunits of Pol II and participate in both transcription and degradation of certain mRNAs [4, 32–34]. All these factors share two common features: they play a role in both transcription and mRNA degradation and can shuttle between nucleus and cytoplasm to transmit information along the mRNA metabolic pipeline. These features are necessary to connect transcription and degradation in mRNA buffering.

Despite extensive knowledge of molecular details that may contribute to mRNA buffering, its mechanism remains mysterious. In this work, we provide the first mechanistic model of mRNA buffering (as far as we realize) that can explain and unify multiple experimental observations of mRNA buffering. In this work, we mainly study the total concentration of all mRNAs, which we call the mRNA concentration in the following unless otherwise mentioned. Our conclusions regarding mRNA buffering do not necessarily apply to all individual genes since they can be under specific regulations [8, 9, 17, 35, 36]. Although most genes exhibit constant mRNA and protein concentration as the cell volume increases, we have shown before that genes with strong (weak) promoters may exhibit sublinear (superlinear) volume-scaling of mRNA and protein concentra-tions [24]. In the following, we show that models involving mRNA feedback to its production or degradation cannot achieve robust mRNA buffering. We then introduce the minimal model of mRNA buffering, the releasing-shuttling (RS) model. The critical ingredients of the RS model are two proteins, X and Y, released after each transcription initiation. Protein X, responsible for mRNA degradation, is exported to the cytoplasm and shuttles back to the nucleus. Protein Y is responsible for the export of mRNA to the cytoplasm. We demonstrate how the RS model quantitatively explains multiple experimental observations and makes experimentally testable predictions. Finally, we identify candidates for the essential proteins X and Y and discuss possible extensions of the RS model.

## RESULTS

### The releasing-shuttling (RS) model

It has been suggested that a negative feedback of mRNA to its own production or a positive feedback to its own degradation can be potential mechanisms of mRNA buffering [14, 17, 26]. However, we found that the feed-back models generally cannot lead to robust buffering. Changing the mRNA production or degradation rates always leads to a significant change in the mRNA concentration (Supplementary Information and Figure S1). In contrast, the mRNA concentration was largely invariant to these perturbations in experiments [8, 9]. In particular, the mRNA concentration and production rate appeared uncorrelated across genome-wide perturbations in mammalian cells [17].

In the following, we introduce a minimal model of mRNA buffering with two essential proteins, X and Y (Figure 1). X is a degradation factor that shuttles between the nucleus and the cytoplasm. Y is an exportation factor and is localized in the nucleus. X can be in three states: the transcription-factor (TF) state in the nucleus that is part of the preinitiation complex (PIC), *X*_*p*_; the released state separated from the PIC right after transcription initiation and ready to be exported to the cytoplasm, *X*_*n*_; the decay-factor (DF) state in the cytoplasm responsible for mRNA degradation, *X*_*c*_. Y can be in two states: the transcription-factor (TF) state in the nucleus that is part of the preinitiation complex (PIC), *Y*_*p*_; the export-factor (EF) state released from the PIC right after transcription initiation and ready to export nuclear mRNAs to the cytoplasm, *Y*_*n*_. We assume that the DF state *X*_*c*_ has a finite rate to shuttle back to the nucleus and become the TF state *X*_*p*_, supported by multiple pieces of evidence showing that mRNA decay factors shuttle in and out of the nucleus and appear critical for transcription [8, 9, 30–32, 37]. Similarly, we assume the EF state *Y*_*n*_ has a finite rate to transition back to the TF state *Y*_*p*_.

**FIG. 1.**
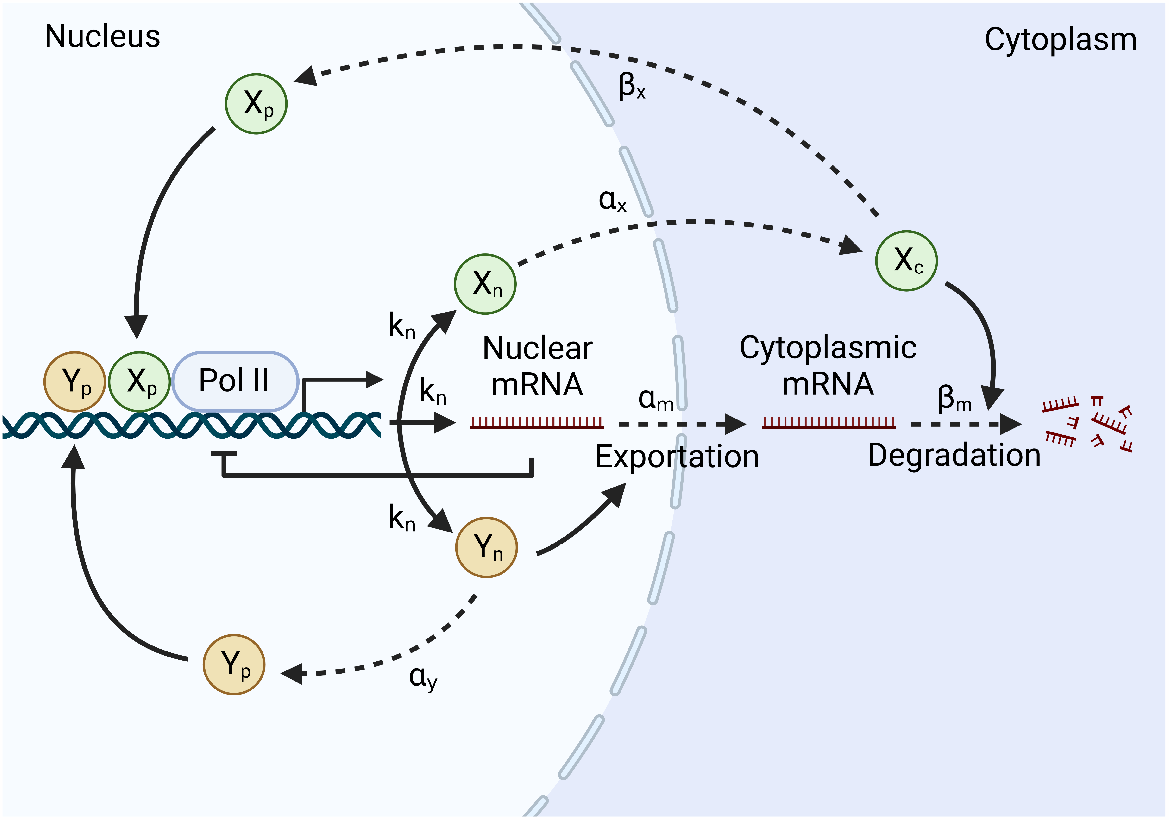
Schematic of the releasing-shuttling model. One copy of protein X and Y in the released states (*X*_*n*_ and *Y*_*n*_) are released after each transcription initiation. *k*_*n*_ is the mRNA production rate, which is also the rate of transcription initiation. *X*_*n*_ transitions to the cytoplasm with a constant rate *α*_*x*_ and becomes *X*_*c*_ in the DF state, which degrades mRNAs with a rate proportional to the constant *β*_*m*_. *X*_*c*_ has a constant rate *β*_*x*_ to shuttle back to the nucleus and becomes *X*_*p*_ in the TF state. *Y*_*n*_ exports nuclear mRNAs to the cytoplasm with a rate proportional to the constant *α*_*m*_. *Y*_*n*_ transitions to *Y*_*p*_ in the TF state with a constant rate *α*_*y*_.

The release of proteins X and Y from the PIC is justified by the fact that transcription initiation is a stepby-step process propelled by different functional groups, which sequentially bind and leave the transcription machinery. In particular, the carboxyl-terminal domain (CTD) region of Pol II is a versatile harbor providing numerous docking points for proteins functioning in diverse processes such as initiation-to-elongation transformation and mRNA processing [38]. We suppose proteins X and Y are two of the signaling molecules that are released during transcription initiation. Recent studies show that mRNA export is under sophisticated regulations [39, 40]. Interestingly, transcription is necessary for mRNA export [41], supporting our model assumption. We hypothesize that protein Y in the state *Y*_*n*_ is an export regulator of mRNA.

The mathematical equations of the RS model are the following,

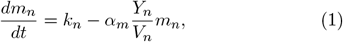

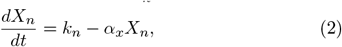

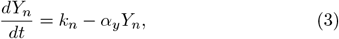

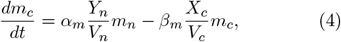

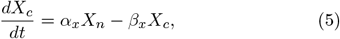

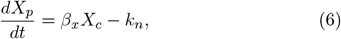

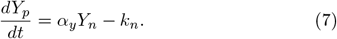

Here *m*_*n*_ is the number of nuclear mRNAs, and *m*_*c*_ is the number of cytoplasmic mRNAs. *k*_*n*_ is the mRNA production rate, which depends on multiple factors, including the concentrations of Pol II, *X*_*p*_, and *Y*_*p*_. *V*_*c*_ is the cytoplasmic volume, and *V*_*n*_ is the nuclear volume that is proportional to *V*_*c*_ [42]. The binding rate of mRNA to the EF state *Y*_*n*_ is proportional to its concentration with a factor *α*_*m*_, which is the limiting step of mRNA export. *α*_*x*_ quantifies how fast *X*_*n*_ escapes the nucleus, and *α*_*y*_ quantifies how fast *Y*_*n*_ transforms back to *Y*_*p*_. Similarly, the binding rate of mRNA to the DF state *X*_*c*_ is proportional to its concentration with a factor *β*_*m*_. *β*_*x*_ quantifies how fast *X*_*c*_ shuttles back to the nucleus.

It is straightforward to find the steady-state solution of the RS model:

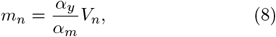

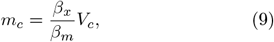

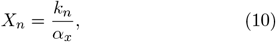

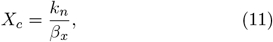

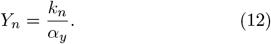

Intriguingly, the numbers of nuclear and cytoplasmic mRNAs are proportional to the nuclear and cytoplasmic volumes, respectively. These are necessary conditions for any model to be biologically valid: the model must predict a constant mRNA concentration. Surprisingly, both the nuclear and cytoplasmic mRNA concentrations are independent of the mRNA production rate *k*_*n*_: the RS model successfully explains the robust mRNA concentration homeostasis against the change of mRNA production rate. This is because the information on the mRNA production rate is conveyed through the amounts of *Y*_*n*_ and *X*_*c*_, synchronizing the speed of mRNA exportation and degradation. To ensure the number of DF state X and EF state Y to be proportional to *k*_*n*_ (Eqs. 11, 12), we concluded that proteins X and Y must be necessary for transcription initiation; otherwise, mRNA buffering is invalid. To explicitly demonstrate this point, we introduced a modified model in which proteins X and Y are not necessary factors to initiate transcription and found that mRNA buffering in this modified model does not hold (Supplementary Information and Figure S2).

Furthermore, it is conceivable that X and Y may also be released at various points throughout the transcription process as research indicates that Ccr4-Not and Xrn1 exert control over not only transcription initiation but also transcription elongation stages [28, 43]. We remark that the exact releasing timings of proteins X and Y are not critical to the predictions of the RS model. Also, the nuclear localization of protein Y is not essential for our conclusions. In a modified model in which Y is exported out of the nucleus and then shuttles back, mRNA buffering is still valid (Supplementary Information and Figure S3).

### The RS model predicts mRNA buffering against various genetic perturbations

The RS model shows that the validity of mRNA buffer-ing does not require a particular form of the mRNA production rate *k*_*n*_ since it does not enter the expressions of *m*_*n*_ and *m*_*c*_. Nevertheless, starting with a biologically valid model for transcription makes it possible to compare theories with experiments explicitly, as we show in the following. Therefore, we incorporated several general features of transcription regulation into the RS model. First, the copy number of Pol II is often limiting for transcription, supported by multiple experiments [20, 21, 26]. Because the Pol II copy number is proportional to the nuclear volume [26], we used the nuclear volume as a proxy for Pol II. It has been found that the number of actively transcribing Pol II increases sublinearly with the total Pol II number given fixed gene copy numbers [26], presumably due to the finite DNA substrates for Pol II to bind [24]. Therefore, we modeled the effects of limiting Pol II resource through an mRNA production rate that is a Hill-function of the nuclear volume, which saturates for a large nuclear volume given a fixed amount of DNA. Second, the nuclear mRNAs have negative feedback to their own production [17, 44] although the detailed mechanisms are still unclear. To sum up, we proposed the following form of the mRNA production rate,

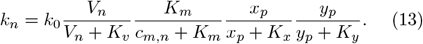

Here, the prefactor *k*_0_ is a constant and the first term on the right side of Eq. 13 represents the sublinear volume dependence of actively transcribing Pol II. The second term represents the negative feedback of nuclear mRNA to transcription where *c*_*m,n*_ = *m*_*n*_*/V*_*n*_. Finally, the last two terms represent the binding probabilities of the transcription factors *X*_*p*_ and *Y*_*p*_ to the PIC. Here, *x*_*p*_ and *y*_*p*_ are the nuclear concentrations of *X*_*p*_ and *Y*_*p*_ respectively: *x*_*p*_ = *X*_*p*_*/V*_*n*_ and *y*_*p*_ = *Y*_*p*_*/V*_*n*_. In the steady state, given the total copy number of protein X (*X*_*t*_) and the total copy number of protein Y (*Y*_*t*_), *X*_*n*_, *X*_*p*_, *X*_*c*_, *Y*_*n*_ and *Y*_*p*_ can be found through the conservation equations, *X*_*t*_ = *X*_*n*_ + *X*_*c*_ + *X*_*p*_, *Y*_*t*_ = *Y*_*n*_ + *Y*_*p*_. Here, *X*_*n*_ = *k*_*n*_*/α*_*x*_, *X*_*c*_ = *k*_*n*_*/β*_*x*_, *Y*_*n*_ = *k*_*n*_*/α*_*y*_, and *k*_*n*_ is a function of *X*_*p*_ and *Y*_*p*_ via Eq. 13.

To mimic genome-wide genetic perturbations similar to previous experiments [8, 9, 17], we systematically perturbed the parameters in Eq. 13 of the RS model, as well as the total copy numbers of proteins X and Y (see details of numerical calculations in Methods). According to the RS model, all the perturbations may change the mRNA production rate but should leave the nuclear and cytoplasmic mRNA concentrations invariant. Indeed, we found that the simulated data nicely matched the experimental data (Figure 2a, b, d, and e) [17], both exhibiting a virtually zero correlation between the mRNA production rate and the mRNA concentrations. In Figure 2a, b, d, and e, the blue dashed lines are *x* = 1 lines, while the red dashed lines are from solving the RS model numerically with different *α*_*m*_, which we explain in the next section. In Ref. [9], Sun et al. measured the mRNA production rate and degradation rate and found an approximately linear relation between these two rates across 46 yeast deletion strains, which suggested mRNA buffering (Figure 2c). To further verify the RS model, we plotted the mRNA production rate vs. the mRNA degradation rate from the same simulations as Figure 2d, e, and also found a linear relationship between them (Figure 2f) in concert with experiments.

**FIG. 2.**
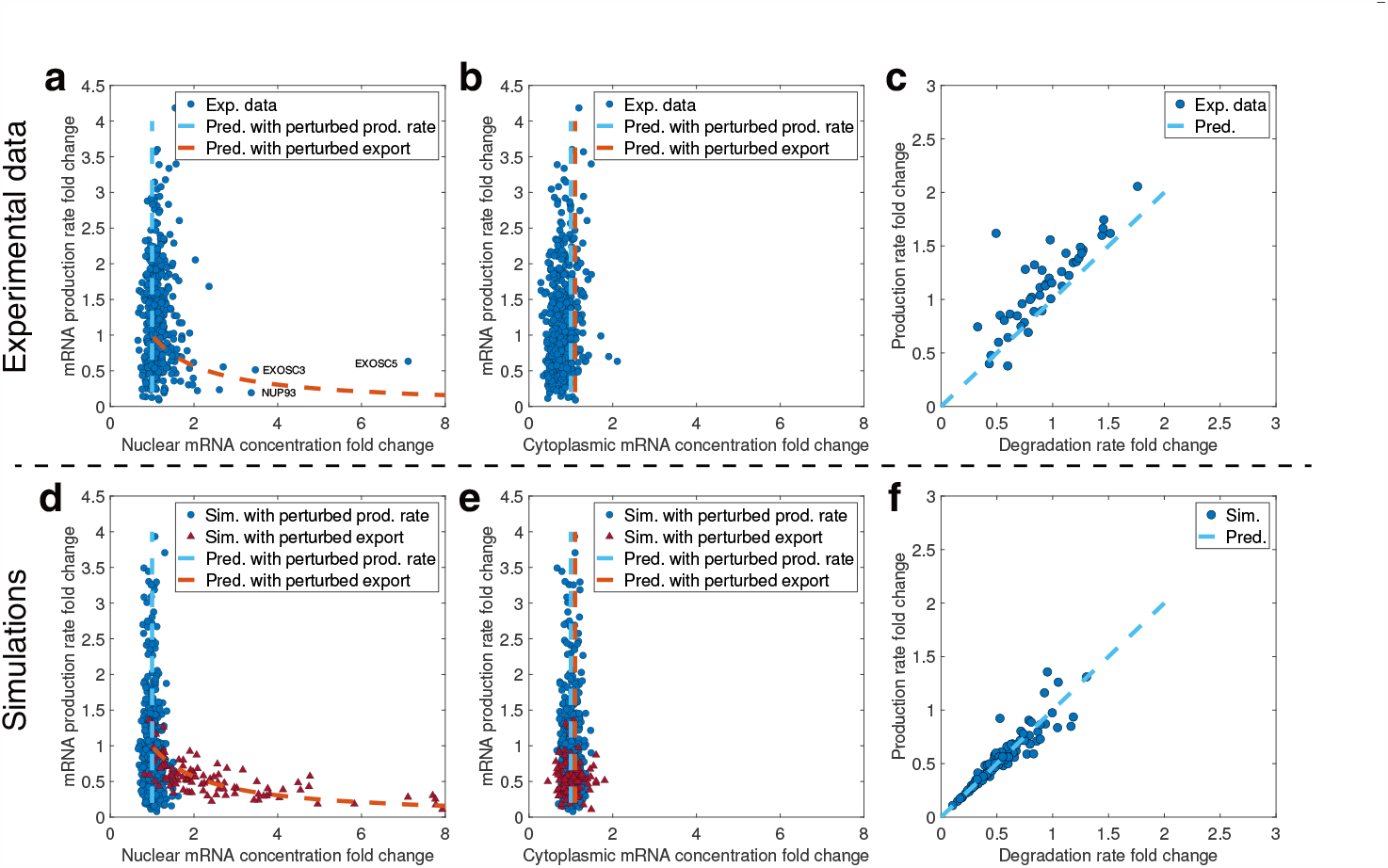
The RS model predicts mRNA buffering against genetic perturbations. Exp., experiment; pred., prediction; prod., production; sim., simulation. (a) Experimental data from Ref. [17] show that the nuclear mRNA concentration is buffered against changes in the mRNA production rate. Each data point represents a knockdown of a particular gene with the three genes EXOSC3, EXOSC5, and NUP93 highlighted. The blue dashed line is an *x* = 1 line, and the red dashed line represents the prediction when nuclear mRNA export is impaired, corresponding to perturbing *α*_*m*_ in the RS model. The same meanings of dashed lines apply to (b, d, e). (b) Experimental data from Ref. [17] show that the cytoplasmic mRNA concentration is buffered against changes in the mRNA production rate. (c) Experimental data of the mRNA production rate vs. the degradation rate from Ref. [9]. The blue dashed line is a *y* = *x* line. (d) Simulations of the RS model show that the nuclear mRNA concentration is buffered against changes in the mRNA production rate. The blue circles represent simulation results with multiple parameters perturbed, and the red triangles represent simulation results with *α*_*m*_ perturbed. The same meanings of points apply to (e). Here, we randomly sampled the mRNA copy numbers from a Poisson distribution with the means equal to the predictions of the RS model, mimicking gene expression noise. We also added Gaussian noise on top of the mRNA production rates so that the CVs equal 0.05. The same noises were applied to (e). (e) Numerical data of the RS model show that the cytoplasmic mRNA concentration is buffered against changes in the mRNA production rate and perturbations in the mRNA nuclear export pathway. (f) Numerical data of the mRNA production rate vs. the degradation rate from the RS model. The blue dashed line is a *y* = *x* line. Data are from the same simulations of (d, e). For both the production and degradation rates, we added Gaussian noises on top of them so that the CVs equal 0.05.

In this work, we used a set of basic parameters (Table II) for all simulations unless otherwise mentioned (Methods). The basic parameters were from the fitting of kinetic data of budding yeast (Figure 4a) with the MM constant *K*_*v*_ inferred from the experimental data of mRNA production rate vs. cell volume (Figure 3b), which we explain in later sections. These parameters are presumably different from mammalian cells. Nevertheless, using the set of basic parameters and only adjusting the *k*_0_ value in Eq. 13, we found that the simulations and experimental data already quantitatively agreed (Figure 2). Therefore, our conclusions regarding the robustness of mRNA buffering are insensitive to the parameters.

**FIG. 3.**
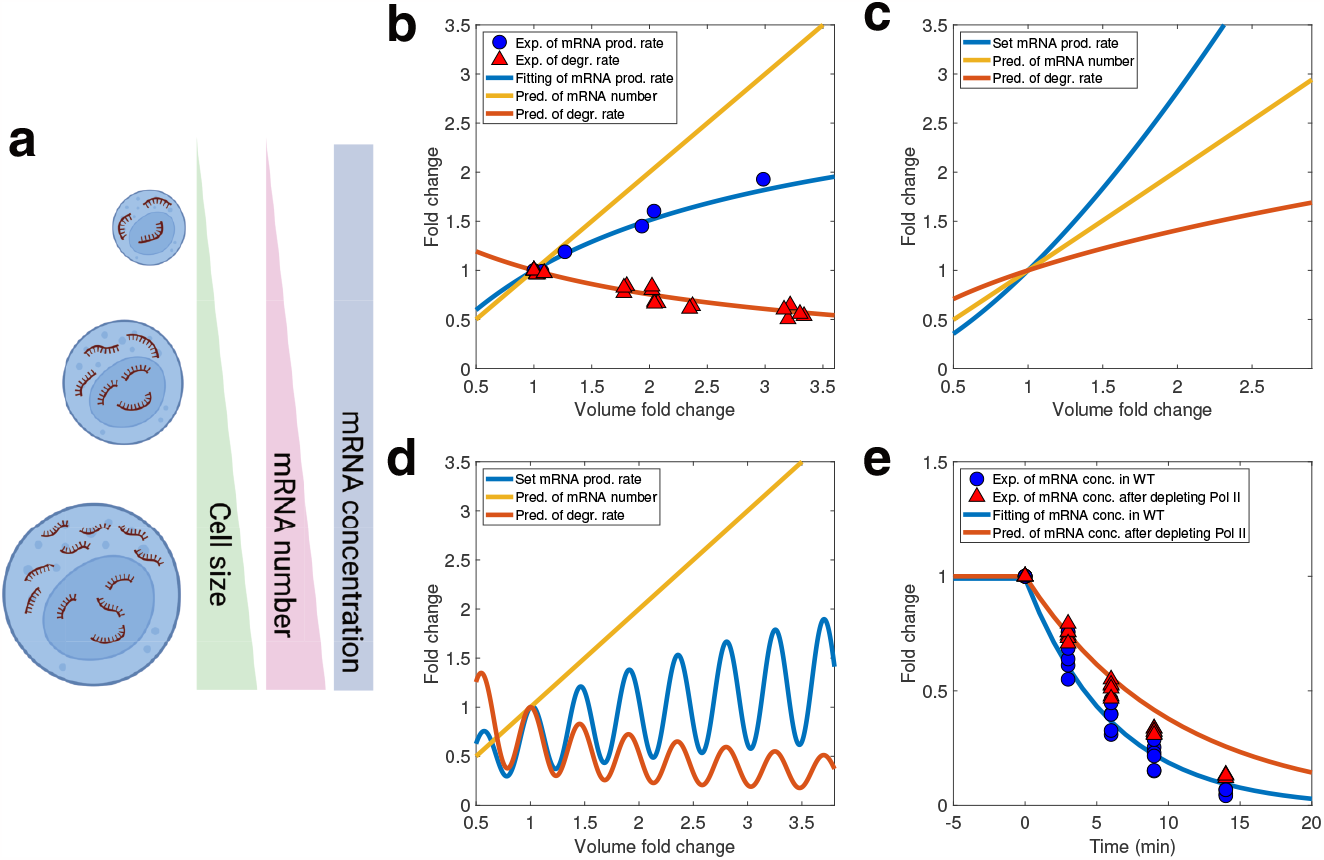
The RS model predicts mRNA buffering during cell growth and slowed mRNA degradation after Pol II depletion. Exp., experiment; degr., degradation; pred., prediction; conc., concentration. (a) Schematic showing the homeostasis of mRNA concentration during cell growth. (b) Experimental data from Ref. [26] for the mRNA production rate, degradation rate, and the mRNA number as a function of cell volume during cell growth. The blue line is a fitting of the experimental data (*r*^2^ = 0.98). The red line is the predicted mRNA degradation rate per mRNA without any fitting parameters from the RS model. The yellow line is the predicted mRNA copy number. (c) mRNA buffering occurs even if the mRNA production rate exhibits a superlinear scaling with cell volume. In this case, the mRNA degradation rate increases with the cell volume accordingly to make the mRNA concentration constant. (d) mRNA buffering occurs even if the mRNA production rate oscillates with cell volume. In this case, the mRNA degradation rate also oscillates, and the mRNA concentration remains constant. (e) The normalized mRNA concentration for a particular gene following transcription shut-off. The blue circles are for cells without Pol II depletion, and the red triangles are for cells with half Pol II depleted. The experimental data are from Ref. [26]. The blue line is a fitting of the experimental data by the RS model (*E*^2^ = 3.87 *×* 10^*−*11^). The red line is the prediction from the RS model for cells with half Pol II depleted without further fitting.

**FIG. 4.**
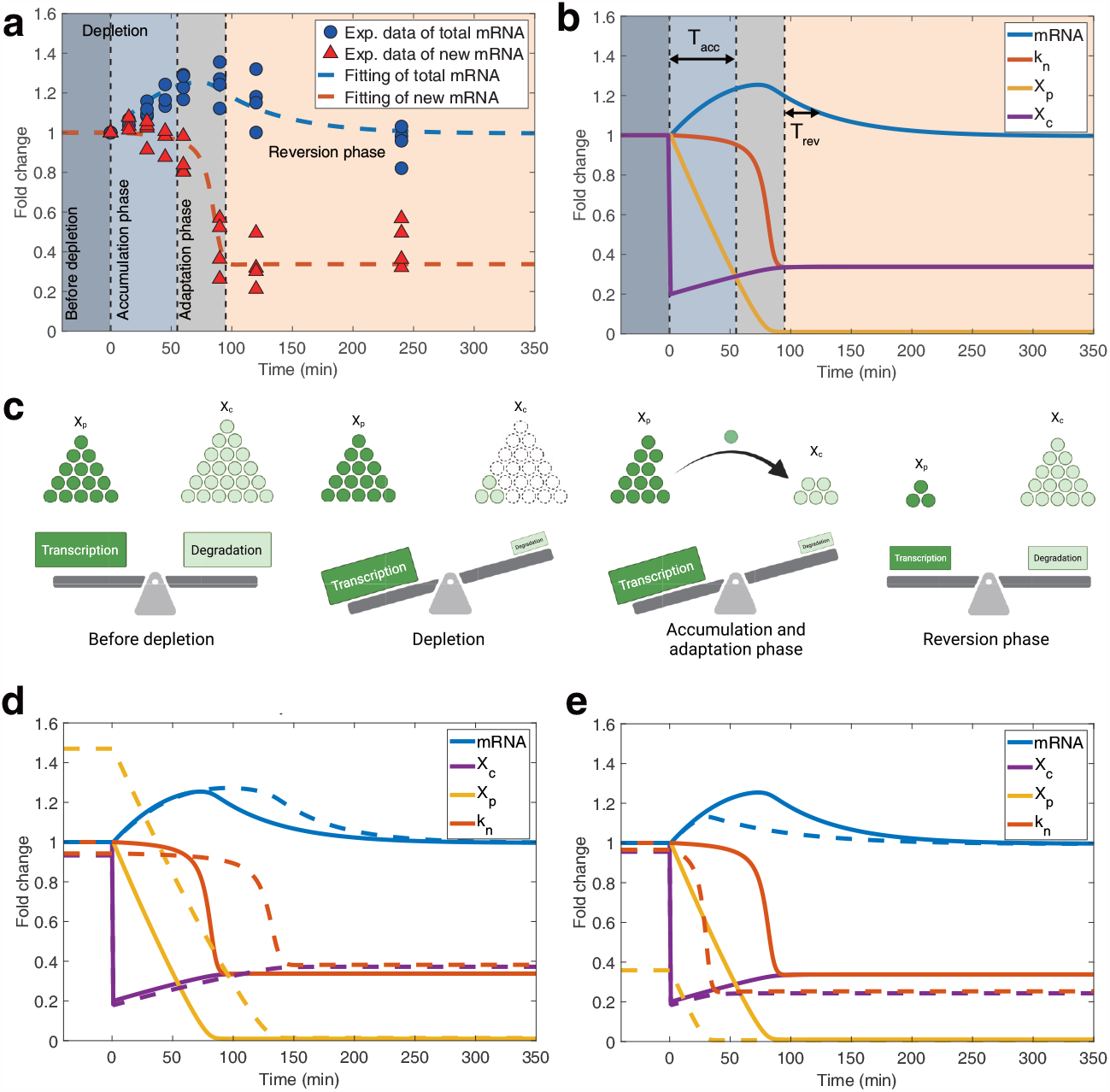
The RS model predicts the temporal change of mRNA concentration after an acute perturbation of the degradation factor. Exp., experiment. (a) The temporal changes in the total mRNA concentration (total mRNA) and recently-transcribed mRNA concentration (new mRNA) following acute depletion of the degradation factor X. The data points represent experimental measurements from [14]. The dashed lines are fitting of the experimental data by optimizing parameters in the RS model (*MSE* = 2.10 *×* 10^*−*3^). We used the mRNA production rate as a proxy for the recently-transcribed mRNA concentration. (b) Simulation results of the temporal dynamics of mRNA, *k*_*n*_, *X*_*p*_, and *X*_*c*_. (c) The schematic illustrating the dynamics of X after an acute depletion of *X*_*c*_ and its influence on transcription and degradation at different times. (d) Impact of a lower Pol II concentration (a larger *K*_*v*_) on the temporal changes following depletion of *X*_*c*_. The solid lines are WT cells, and the dashed lines are for cells with a larger *K*_*v*_. Both the solid and dashed lines are relative to the values of the WT cells before depletion. (e) Influence of a lower X concentration (a lower *X*_*t*_) on the temporal changes after depleting *X*_*c*_. The lines have the same meaning as (d).

### The RS model predicts when mRNA buffering break downs

Experimentally, violations of nuclear mRNA buffering were observed when the export of nuclear mRNA was impaired [17]. Intriguingly, the RS model also predicts the breakdown of nuclear mRNA buffering. Specifically, the RS model predicts that when nuclear mRNA export is impaired, that is, a lower *α*_*m*_ in the RS model (Eq. 8), the nuclear mRNA concentration is not constant anymore but negatively correlates with the mRNA production rate. In Figure 2a and d, the red dashed lines were from the numerical calculations of the RS model with changing *α*_*m*_. The red triangles in Figure 2d were obtained by randomly sampling *α*_*m*_ (see details of numerical calculations in Methods). In the RS model, the negative correlation between the mRNA production rate and the nuclear mRNA concentration comes from the negative feedback from the nuclear mRNA concentration to the mRNA production rate, and we confirmed this by showing that the negative correlation is absent in a modified model without this negative feedback (Figure S4). Intriguingly, the RS model predicts that the cytoplasmic mRNA concentration is uninfluenced by the impairment of nuclear mRNA export (Eq. 9), which exactly matches the experimental observations (Figure 2b and e).

Biologically, nuclear mRNA export needs accurate mRNA processing and the proper assistance of multiple structures like nuclear pores [45]. The RS model successfully explains the breakdown of nuclear mRNA buffering when the nuclear pore component NUP93 and the core components of the nuclear RNA exosome EXOSC3 and EXOSC5 are knocked down [17], strongly supporting that the RS model captures the key ingredients of mRNA buffering.

### The RS model leads to mRNA buffering as the cell volume increases

We then sought to test if the RS model can achieve homeostasis of mRNA concentration during cell growth (Figure 3a). For simplicity, we assumed that the total copy numbers of proteins X and Y are proportional to the cell volume, which is valid for most proteins [46]. We first fit the experimental data of mRNA production rate vs. cell volume from Ref. [26] using Eq. 13 assuming a constant ratio between the total cell volume and nuclear volume (the blue line in Figure 3b). From this fitting, we obtained the value of the MM parameter *K*_*v*_ for WT cells (Eq. 13). *K*_*v*_ and other parameters inferred from the fitting of kinetic data of budding yeast (see the next section) constitute the basic parameters in this work (Methods and Table II). In the above fitting, we neglected the volume dependence of the concentrations of transcription factors, *x*_*p*_ and *y*_*p*_ and confirmed that this approximation was valid (Figure S5). The nuclear mRNA concentration *c*_*m,n*_ is also constant in Eq. 13 due to mRNA buffering, as shown in the following.

We defined the mRNA degradation rate *δ*_*m*_ as the degradation rate per mRNA, i.e., *δ*_*m*_ = *β*_*m*_*X*_*c*_*/V*_*c*_ according to Eq. 4. Therefore, the mRNA lifetime *τ*_*m*_ = 1*/δ*_*m*_. Notably, the predicted mRNA degradation rate from the RS model (the red line in Figure 3b) precisely matches the experimentally measured values (red triangles) with-out any fitting parameters. Within the RS model, while the mRNA production rate scales sublinearly with the cell volume, the degradation rate decreases accordingly to achieve a constant mRNA concentration (Figure 3b). The RS model shows that the mRNA degradation rate automatically compensates for the sublinear mRNA production rate, consistent with experiments. In fact, according to the RS model, any cell-volume dependence of the mRNA production rate will lead to a constant mRNA concentration. We confirmed this prediction by assuming an mRNA production rate that is a superlinear (Figure 3c) or oscillating (Figure 3d) function of cell volume. In both cases, the mRNA degradation rate self-adjusts to compensate for the mRNA production rate and generates a constant mRNA concentration. This stringent prediction can be tested experimentally.

### Slowed mRNA degradation after Pol II depletion

In Ref. [26], Swaffer et al. depleted half of Pol II in budding yeast and measured the degradation of a particular gene’s mRNA. They found that mRNA degradation slowed down after Pol II depletion. We found that this experimental observation is a direct consequence of the RS model. According to the RS model, the mRNA degradation rate should decrease if the mRNA production rate decreases. Because the Pol II copy number is proportional to the nuclear volume, reducing the Pol II concentration to its half value is equivalent to doubling the MM parameter *K*_*v*_ in Eq. 13. Once the mRNA production rate decreases due to Pol II depletion, the number of the DF state *X*_*c*_ in the cytoplasm decreases according to Eq. 11 since the replenishment of *X*_*c*_ becomes slower. The reduction of *X*_*c*_ leads to slowed mRNA degradation. Due to the conservation of total protein X number, the number of the TF state *X*_*p*_ increases, also alleviating the reduction of mRNA production rate.

To verify our idea, we numerically simulated a slightly modified model in which we separated the transcription of a single gene from the rest of the genome. We monitored the time-dependence of the mRNA number of the particular gene (including both the nuclear and cytoplasmic mRNA) after turning off its transcription. We first fitted the data of mRNA production rate vs. cell volume before Pol II depletion by optimizing parameters except *K*_*v*_ in the set of basic parameters (blue lines and circles in Figure 3e, and Methods). We then doubled *K*_*v*_ to model Pol II depletion. We found that the RS model nicely predicts the slowed mRNA degradation after Pol II depletion without further fitting (red lines and triangles in Figure 3e).

### Temporal transcription dynamics after rapid degradation of protein X

In Ref. [14], Chappleboim et al. rapidly depleted Xrn1 in budding yeast and monitored the temporal dynamics of the total mRNA concentration and the level of recently transcribed mRNA, which is a good proxy for the mRNA production rate. Experimental observations revealed three phases of mRNA dynamics upon the sudden removal of the protein Xrn1 (Figure 4a). In the accumulation phase, the mRNA concentration increased, and the mRNA production rate remained almost constant. In the adaptation phase, the mRNA production rate dropped rapidly, and the mRNA concentration ceased to increase. In the reversion phase, the mRNA production rate reached the new steady-state value, and the mRNA concentration gradually reduced to its original value before the depletion of Xrn1. To verify whether the RS model can explain the temporal dynamics of mRNA as a more stringent test, we studied the RS model after a rapid depletion of protein X.

First, we remark that in the RS model, protein X is presumably a coarse-grained combination of several proteins. This idea is supported by the experimental observations that mRNA buffering was still valid even if Xrn1 was knocked out [8, 14], which we discuss in detail later. Therefore, the depletion of Xrn1 should correspond to a partial depletion of protein X in the RS model. Because Xrn1 is the primary degradation factor in budding yeast [8] and more than 90 percent copies of Xrn1 are localized in the cytoplasm [47], we modeled the depletion of Xrn1 as a rapid reduction of *X*_*c*_ to its 20% value in the cytoplasm. We neglected the depletion of protein X in the nucleus, including the TF state *X*_*p*_ and the released state *X*_*n*_ for simplicity. This assumption can be relaxed as long as the nuclear X does not decrease significantly.

Due to the rapid depletion of the DF state *X*_*c*_ in the cytoplasm, the replenishment of *X*_*p*_ (Eq. 6) significantly decreases (Figure 4b). However, we argue that a delay between the decrease of the mRNA production rate and the reduction of *X*_*p*_ can emerge because the transcription factor concentration is still much higher than the Michaelis-Menten constant *K*_*x*_ (Eq. 13). In this case, the mRNA production rate will be maintained at a constant value for a finite duration, in agreement with experiments [14]. Therefore, the TF state *X*_*p*_ initially decreases linearly because of the conversion from *X*_*p*_ to *X*_*n*_ (Eq. 6). The linear reduction of *X*_*p*_ holds until *X*_*p*_ hits *K*_*x*_, which leads to a significant decrease in the mRNA production rate (Eq. 13) (Figure S6). Therefore, we can approximate the duration of the accumulation phase as *T*_*acc*_ *≈* (*X*_*p*_ *− K*_*x*_)*/*(*k*_*n,bf*_ *f*_*dep*_) according to Eq. 6. Here, *k*_*n,bf*_ is the mRNA production rate before perturbation, which is also the replenishment rate from *X*_*c*_ to *X*_*p*_. We have taken account of the reduced replenishment rate as (1 *− f*_*dep*_)*k*_*n,bf*_ where *f*_*dep*_ is the fraction of depleted *X*_*c*_ (80% in this case). The values of *X*_*c*_, *X*_*p*_, and the mRNA production rate then quickly reach the new steady-state values, after which the mRNA concentration gradually returns to its original value with a relaxation time determined by the mRNA degradation rate, *δ*_*m*_ = *β*_*m*_*X*_*c*_*/V*_*c*_ according to Eq. 4. Therefore, the duration of the reversion phase is *T*_*rev*_ *≈ V*_*c*_*/β*_*m*_*X*_*c*_. Figure 4c shows a schematic of this process.

In the above discussion, the initial value of the transcription factor concentration *x*_*p*_ must be far above the parameter *K*_*x*_ (Figure S6). This condition suggests that for WT cells, *X*_*p*_ is generally non-limiting for transcription, although it is still necessary to initiate transcription. We also confirmed that the three phases in the transcription dynamics following an acute depletion are robust against different choices of the parameters (Figure S7). In conclusion, the predictions of the RS model not only perfectly aligned with the experimental data from which we inferred the set of basic parameters (Figure 4a and Methods) but also provided valuable insights into the underlying mechanism (Figure 4b and c).

Next, we sought to systematically investigate the transcription dynamics after acute depletion of *X*_*c*_ according to the RS model, in particular, considering cells with different concentrations of Pol II or protein X. Future experiments can test our predictions. We compared a WT cell (solid lines in Figure 4d) and a mutant cell with a lower Pol II concentration (dashed lines in Figure 4d). The mutant with a lower Pol II concentration, equivalent to a larger *K*_*v*_ in the RS model (Eq. 13), has a mildly lower mRNA production rate than the WT cell. Therefore, the mutant has fewer *X*_*n*_, *X*_*c*_ and more *X*_*p*_ (Eqs. 10 and 11). Consequently, it takes longer for the mutant cell to reach the adaptation phase since it needs a longer time for the TF state *X*_*p*_ to decrease and reach the MM constant *K*_*x*_. Because of the longer duration of the accumulation phase for the mutant, which also has a similar accumulation rate of *X*_*c*_ as the WT cell, the mutant has a higher *X*_*c*_ in the new steady state. Therefore, the mutant recovers more rapidly during the reversion phase than the WT cell.

For the mutant (dashed lines in Figure 4e) with a lower protein X concentration, it has fewer TF state *X*_*p*_ than the WT cell (solid lines in Figure 4e). However, its mRNA production rate remains close to the WT cell as long as the concentration of *X*_*p*_ is still significantly larger than *K*_*x*_. Because of the fewer *X*_*p*_, the mutant with a lower X concentration exhibits a shorter accumulation phase than the WT cell because the duration of the accumulation phase depends on the initial *X*_*p*_, *T*_*acc*_ *≈* (*X*_*p*_*−K*_*x*_)*/*(*k*_*n,bf*_ *f*_*dep*_). Therefore, the mutant also has a lower number of *X*_*c*_ during the reversion phase, so the mRNA concentration recovers slower in the mutant than in the WT cell. We also investigated the temporal transcription dynamics for other mutants with different modified parameters (Figure S7).

### Candidates of X and Y

Experiments showed that the mRNA concentration was buffered even when the global degradation factors such as Dcp2 and Xrn1 were knocked out [8, 14]. These factors play vital roles in 5’-3’ mRNA degradation, the predominant pathway of mRNA degradation [31]. These observations suggest that the protein X and Y in the RS model may not represent a single protein, and they may be a combination of multiple proteins. To demonstrate this idea explicitly, we investigated the response of mRNA concentration to an abrupt drop in the mRNA production rate by considering cells with different total numbers of protein X. We found that the recovery time of mRNA concentration diverges in the limit of *X*_*t*_ *→*0 (upper panel of Figure 5a). This result suggests that if protein X is only Xrn1, cells with Xrn1 knocked out should not recover from a slight noise in the mRNA production rate, so it cannot achieve mRNA buffering, contradicting experiments. We analyzed protein Y similarly and obtained the same results (lower panel of Figure 5a). Therefore, we proposed that X and Y represent groups of multiple proteins that perform similar functions (Figure 5b).

**FIG. 5.**
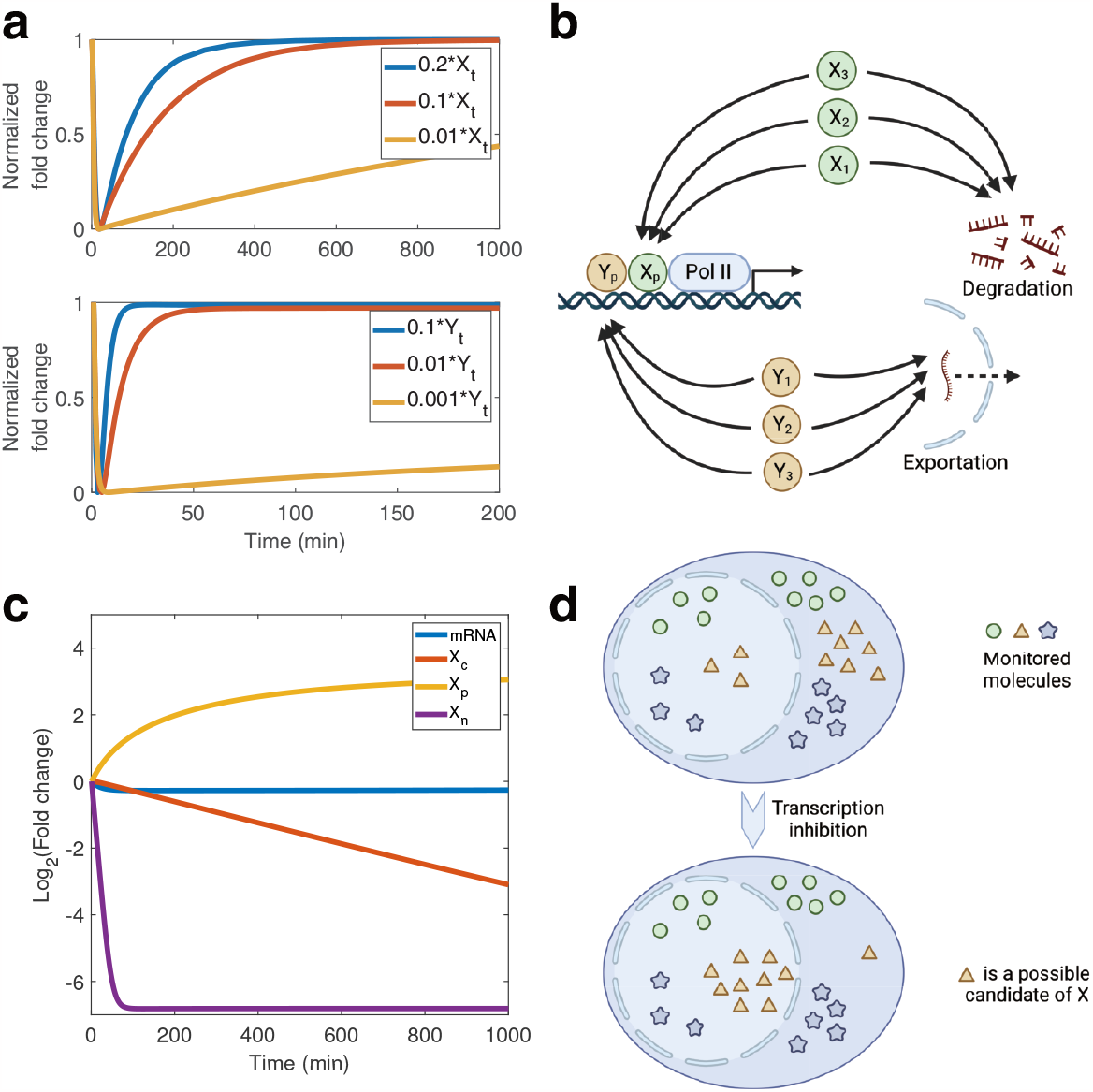
Proteins X and Y should represent groups of proteins with similar functions. (a) The recovery of cytoplasmic mRNA (upper panel with varying copy numbers of X) and nuclear mRNA (lower panel with varying copy numbers of Y) after a rapid perturbation to the mRNA production rate. The y-axis is shifted and normalized so that the value before perturbation is one and the minimum value is 0. (b) Schematic illustrating X and Y can be groups of different proteins executing similar functions. (c) The time course of the mRNA concentration, *X*_*c*_, *X*_*p*_, and *X*_*n*_ after the complete shutoff of transcription. (d) Protein X in the cytoplasm will gradually shuttle to the nucleus until most of its copies are localized in the nucleus after a complete transcription inhibition, which can be used to detect the candidates of protein X.

The necessity of the two transcription factors, *X*_*p*_ and *Y*_*p*_ to transcription (Figure S2) implies that they must be essential components of the transcription machinery, such as the subunit of Pol II, the general transcription factors, and other necessary molecules. Rpb4 and Rpb7, the subunits of Pol II, are possible candidates involved in shuttling across the nuclear membrane and regulating mRNA degradation [4, 32–34]. Although the cytoplasmic function of Rpb4 is shown to be dispensable [48], similar functions may exist in other factors of the transcription machinery. CDK8, orthologous to budding yeast Ssn3, is a cyclin-dependent kinase (CDK) functional in phosphorylating the CTD of Pol II, promoting the assembly of the elongation complex [49]. Evidence shows that CDK8 participates in the degradation of several mRNAs related to metabolism [50, 51]. Hmt1 is a methyltransferase in budding yeast, which participates at the beginning of the transcriptional elongation process and influences mRNA exportation by methylating some RNA-binding proteins (RBPs) [52]. SUS1 encodes a protein that is a shared subunit of two complexes: Spt-Ada-Gcn5 acetyltransferase (SAGA) and transcription and export complex 2 (TREX2). SAGA modifies chromatin structure and facilitates transcription initiation, while TREX-2 is a complex that associates with the nuclear pore complex and mediates mRNA export [53]. In a nutshell, we provided a list of candidates of X and Y, participating in at least two of the three processes, including transcription, exportation, and degradation (Table I).

**TABLE 1.** Candidates of X and Y in *S. cerevisiae*.

| Gene name | Functions as | Gene name | Functions as | Gene name | Functions as | Gene name | Functions as |
| --- | --- | --- | --- | --- | --- | --- | --- |
| CAF40 | X | MOT2 | X | NUP84 | Y | SPN1 | Y |
| CCR4 | X | NAB2 | X and Y | NUP85 | Y | SPT6 | Y |
| CDC36 | X | NAP1 | Y | PHO85 | Y | SSL2 | Y |
| CDC39 | X | NOT3 | X | POP2 | X | SSN3 | X |
| CLA4 | Y | NOT5 | X | RAT1 | X | SUS1 | X and Y |
| DCP2 | X | NUP100 | Y | RMD9 | X | THO2 | Y |
| HMT1 | Y | NUP120 | Y | RPB4 | X and Y | UTP8 | Y |
| HPR1 | Y | NUP133 | Y | RPB7 | X | UTP9 | Y |
| HTZ1 | X | NUP145 | Y | RSP5 | Y | WTM1 | Y |
| ISW1 | Y | NUP159 | Y | RTG2 | Y | XRN1 | X |
| MGA2 | X | NUP42 | Y | SEC13 | Y | YAR1 | Y |
| MLP2 | Y | NUP53 | Y | SGF73 | Y | YGR122W | X |

Typically, genetic perturbation methods map a gene to a phenotype of interest. If the knockout or knockdown of a particular gene affects the phenotype, it suggests that the gene is likely responsible for the phenotype. However, in the case of protein X, presumably a combination of proteins with redundant functions, perturbing a single gene may not significantly impact mRNA buffering, making it challenging to identify the candidates responsible for mRNA buffering using traditional approaches. Fortunately, the RS model suggests an alternative solution. We found that the RS model made an interesting prediction if transcription is completely inhibited, e.g., through rapid Pol II depletion or transcription-inhibiting drugs. In this case, the transformation from the TF state *X*_*p*_ to the released state *X*_*n*_ is blocked and *X*_*n*_ quickly reduces to zero. Meanwhile, the DF state *X*_*c*_ in the cytoplasm gradually shuttles back to the nucleus to become *X*_*p*_ (Figure 5c). During this process, mRNA continues to be degraded and eventually stabilizes at a lower level (Figure 5c). By monitoring the cytoplasmic proteins that are predominantly transported back to the nucleus after transcription shut-off, one can identify the possible constituents of X (Figure 5d). This prediction of the RS model provides an experimental protocol to examine the potential candidates for X.

### Extensions of the RS model

The RS model provides a fundamental understanding of transcription regulation by revealing the coordination of mRNA production, exportation, and degradation rates to maintain a constant mRNA concentration. However, it is essential to recognize that the real-world dynamics of mRNA regulation can be more complex than the current RS model. This section discusses some of the complexities that may go beyond the simplified scenario of mRNA buffering and how the RS model can be extended to address them.

We first discuss the mRNA dynamics during viral infection. When cells are infected, the mRNA degradation rates are enhanced to facilitate the rapid turnover of host mRNA and redirect cellular resources toward viral replication, a phenomenon called “host shutoff” [54]. Contrary to mRNA buffering, the mRNA production rate becomes slower, although the mRNA degradation rate is accelerated by viral infection [55]. Recently, Gilbertson et al. showed that in mammalian cells, the poly(A)-binding protein cytoplasmic 1 (PABPC1) plays a critical role in this process [37]. PABPC1 is an RBP that binds directly to the poly(A) tail of the pre-mRNA in the nucleus and is released in the cytoplasm after mRNA is degraded, functioning in mRNA translation and degradation [56–58]. Researchers found that it also interferes with the formation of PIC in the nucleus [37]. Once the viral endonuclease accelerates mRNA degradation, more PABPC1 proteins are released into the cytoplasm, which then shuttle to the nucleus and inhibit transcription [37].

With this experimental evidence, we modified the RS model by adding a third protein P representing PABPC1 (Figure 6a and Methods). In this modified model, there are four states of P, including the one binding to the nuclear mRNA (*P*_*n,b*_), the one binding to the cytoplasmic mRNA (*P*_*c,b*_), the free P in the nucleus (*P*_*n*_), and the free P in the cytoplasm (*P*_*c*_). We assumed that protein P binds mRNA tightly and gets released only when the mRNA is degraded so that the numbers of *P*_*n,b*_ and *P*_*c,b*_ are equal to the nuclear mRNA and cytoplasmic mRNA, respectively. *P*_*n,b*_ is assumed to be exported along with mRNA, while the rate of *P*_*c*_ shuttling into the nucleus is determined by the factor *β*_*p*_. We also modeled the inhibitions of *P*_*n*_ to the mRNA production rate as a Hill function. We modeled the effects of viral infection as an increased mRNA degradation rate by increasing *β*_*m*_ (Eq. 4). Simulations showed that upon viral infection, the mRNA degradation rate per mRNA increases while the mRNA production rate decreases so that the mRNA concentration decreases (Figure 6b, and see Methods for the details of the simulations). In the meantime, protein P is enriched in the nucleus, in agreement with experimental observations during viral infection.

**FIG. 6.**
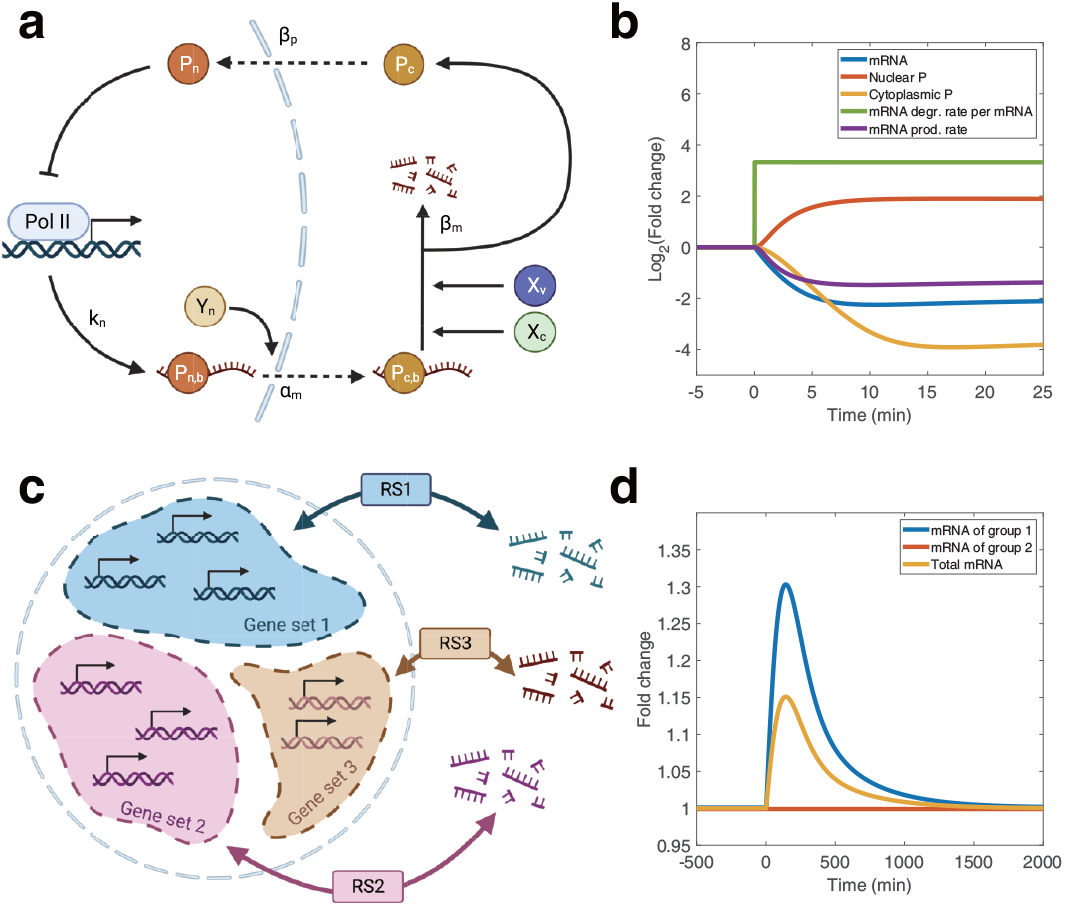
The extensions of the RS model. Degr., degradation; prod., production. (a) Schematic of the modified model in which a third protein P is added. *X*_*v*_ represents the viral endonuclease that increases the factor *β*_*m*_. (b) Simulations of the dynamics of the total mRNA concentration, nuclear P (*P*_*n*_ + *P*_*n,b*_), cytoplasmic P (*P*_*c*_ + *P*_*c,b*_), degradation rate per mRNA, and production rate after a sudden increment of *β*_*m*_ at time 0. (c) Schematic of the generalized model in which different sets of genes are regulated by different groups of X and Y, e.g., RS1, to buffer mRNA concentration separately. (d) Simulations of the dynamics of two groups of mRNA regulated by distinct groups of X and Y and their sum after a sudden decrement in *X*_*c*_ of group 1 at time 0.

Next, we discuss the differential regulations of specific mRNA groups [14, 29, 31, 59]. Recently, Chattopadhyay et al. showed that blocking the import of Xrn1 to the nucleus significantly reduces the production and degradation rate of a subset of mRNAs but does not affect the rest [29]. Another example is that the expression of ribosomal genes responds differentially to the knock-out of Ccr4-Not and Xrn1 [59]. Interestingly, in both cases, mRNAs related to ribosome biogenesis appear to be regulated distinctly. Given these experimental observations, we propose that separate sets of proteins X and Y can coexist to buffer each group of mRNA independently (Figure 6c), contributing to the overall homeostasis of mRNA concentration. One benefit of separate sets of proteins X and Y is that the expression changes of one group of genes do not significantly influence the other groups. To demonstrate this benefit explicitly, we modified the RS model by introducing two sets of X and Y, regulating two groups of genes. Simulations of this modified RS model show that despite changes in the mRNA concentration in one group after perturbing its *X*_*c*_, the mRNA concentration of the other group is largely invariant (Figure 6d, see Methods for the details of the simulations).

In summary, while the RS model provides a powerful framework for understanding mRNA buffering, its simplicity also allows for straightforward extensions and adaptations to capture the complexity of mRNA regulation. By incorporating additional regulatory information, the RS model can be tailored to address specific biological contexts and shed light on the diverse regula-tion mechanisms of the mRNA level.

## DISCUSSIONS

In this study, we introduced the releasing-shuttling model as a minimal mechanistic framework to explain mRNA buffering, a fascinating phenomenon observed in various organisms where mRNA concentrations remain approximately constant through coupled changes in mRNA production and degradation rates. The RS model incorporates two essential proteins, X and Y, involved in transcription, exportation, and degradation. The model demonstrates that the clue of the mRNA production rate is conveyed downstream via X and Y, synchronizing all rates to achieve mRNA buffering. Through mathematical modeling and analysis, we have shown that the RS model provides a robust mechanism of mRNA buffering and makes valuable predictions that are experimentally testable. The RS model also predicts mRNA buffering as the cell volume grows, no matter how the mRNA production rate depends on cell volume. Meanwhile, we noticed that the concentrations of mRNA and protein decrease in budding yeast cells with extremely large sizes [60], which can be due to the imbalance of biomass production and osmolyte synthesis [61, 62]. The RS model also quantitatively predicts the mRNA dynamics following the rapid depletion of protein X. In particular, the RS model predicts a rapid drop in mRNA degradation rate but a relatively constant mRNA production rate in the accumulation phase followed by the adaptation phase and then the reversion phase in which the mRNA concentration decreases exponentially.

An intriguing aspect of the RS model is the possibil-ity that proteins X and Y may be groups of molecules with redundant functions, providing a plausible explanation for the robustness of mRNA buffering in the face of individual genetic perturbations. It is important to note that the precise identity of proteins X and Y in different organisms and cell types remains an open question. We provided a list of candidates for these proteins in budding yeast, including subunits of RNA polymerase II (Pol II), general transcription factors, and proteins involved in mRNA export and degradation (Table I). Specifically for X, we provided an experimentally feasible method to examine the candidate. Future experiments can utilize the predictions of the RS model to identify the specific proteins responsible for mRNA buffering in different cellular contexts. The RS model also shows flexibility for extensions to capture complexities in biological systems. By incorporating an additional protein, the RS model successfully predicts opposite changes in the mRNA production rate and degradation rate during the viral infection, contrary to mRNA buffering. Furthermore, separate sets of proteins X and Y can coexist and regulate different mRNA subsets, highlighting the versatility of the RS model in capturing complex regulatory scenarios.

While the RS model successfully explains mRNA buffering and makes valuable predictions, it also raises important questions and opportunities for further investigation. In Ref. [29], the authors mentioned that the total mRNA concentration is unaffected by mutations in the NLS of Xrn1. However, the RS model predicts a lower mRNA concentration if Xrn1, as a global decay factor of mRNA, is retained in the cytoplasm, which mathematically can be considered as a lower *β*_*x*_ value in Eq. 9. We also noticed that the RS model does not require a persistent binding between mRNAs and proteins X and Y. In other words, we found that mRNA imprinting of proteins X and Y does not appear necessary to achieve mRNA buffering [34]. Phenomenological models of mRNA imprinting have been built in which the crosstalk factors impact the transcription and mRNA stability [36]. In summary, further quantitative data and theoretical analysis are needed to guide the modification of the RS model to incorporate other important biological features.

## METHODS

### The modified model for the viral infection

To explain the experimental observations of the viral infection, we added another protein, P, to the RS model, representing protein PABPC1 (Figure 6a). The dynamics of P follows

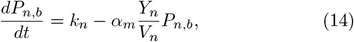

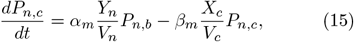

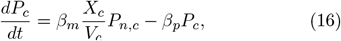

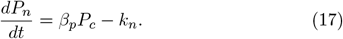

Here *P*_*n,b*_ = *m*_*n*_ and *P*_*c,b*_ = *m*_*c*_ because P binds to mRNA tightly. To model the inhibition of PABPC1 to transcription, we included the free protein P, *P*_*n*_, to Eq. 13, so that

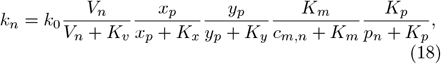

where *p*_*n*_ is the concentration of *P*_*n*_ in nucleus, and *K*_*p*_ is a constant.

#### Analysis of experimental data

In Figure 2a, b, the experimental data were obtained from [17]. The nuclear mRNA concentrations, cytoplasmic mRNA concentrations, and mRNA production rates were calculated by averaging the “mean nucleus mean FISH,” “mean cytoplasm mean FISH,” and “mean nucleus mean EU” values, respectively, across all samples perturbing the same gene (or scrambled pieces). The data were then normalized by the mean values of the scrambled samples to obtain fold changes.

In Figure 3e, the experimental data for WT cells and cells with half of the Pol II depleted were obtained from Supplementary Figure 8e of [26]. We set the time point at 6 minutes as the starting time point, the values of which were set to 1. Only the data after the starting time point were used. We then fitted the data of WT cells with a one-phase exponential decay model to get the degradation rates per mRNA (*r*^2^ = 0.97).

In Figure 4a, the experimental data were obtained from the supplementary table “xrn1 global nas traj repeats” in [14]. mRNA levels were obtained from “smoothed mRNA”, and newly transcribed mRNA levels were obtained from “nascent”. All values were normalized with the corresponding values at time 0 to obtain fold changes.

#### Determination of basic parameters

In the RS model, there are a total of 14 unknown parameters: *k*_0_, *K*_*x*_, *K*_*y*_, *K*_*m*_, *K*_*v*_, *α*_*m*_, *α*_*x*_, *α*_*y*_, *β*_*m*_, *β*_*x*_, *X*_*t*_, *Y*_*t*_, *V*_*n*_, and *V*_*c*_. Exploring the entire parameter space to find combinations that quantitatively fit the experimental data is a complex task due to the large number of parameters involved. Therefore, we fixed some parameters within biologically reasonable ranges to simplify the analysis and made some reasonable assumptions. More quantitative experiments are necessary to refine the parameters and improve our understanding of the RS model in the future.

We assumed a total cell volume of 40 fL, the typical volume of haploid *S. Saccharomyces*. Since the nuclear volume is approximately 7% of the total cell volume [63], we set the nuclear volume (*V*_*n*_) to 2.8 fL and the cytoplasmic volume (*V*_*c*_) to 37.2 fL. Throughout the simulations, the values of *V*_*n*_ and *V*_*c*_ are kept fixed, except for the simulations involving cell growth (Figure 3b, c, and d), where the ratio between *V*_*n*_ and *V*_*c*_ is maintained while the cell volume increases.

To estimate the value of *K*_*v*_, we assumed that the influence on the mRNA production rate *k*_*n*_ (Eq. 13) is primarily determined by the term *V*_*n*_*/*(*V*_*n*_ + *K*_*v*_) when the cell volume increases, which we confirmed numerically (Figure S5). We calculated the fold changes of mRNA production rates 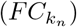 in cells with different volumes as

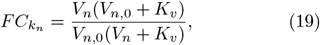

where *V*_*n*,0_ is the reference nuclear volume. We fit this equation to the experimental data from [26] to determine the value of *K*_*v*_ (Figure 3b).

We performed a fitting process to determine the remaining 11 parameters using experimental data from [14]. We searched the optimal fitting parameters starting from the biologically reasonable values inferred from the experimental data. We used the following lower indexes to indicate the values of different factors at different phases of Xrn1 depletion: “bf” (before depletion), “1” (start of the accumulation phase), “acc” (during the accumulation phase), “2” (start of the adaptation phase), “adp” (during the adaptation phase), “3” (start of the reversion phase), and “rev” (during the reversion phase). Figure S8 shows a schematic of the fitting process.

We assumed that *α*_*y*_ is sufficiently large so that *Y*_*p*_ is virtually proportional to *k*_*n*_ at any given time, i.e., *Y*_*p*_ *≈ k*_*n*_*/α*_*y*_, which ensures a constant nuclear mRNA concentration (note that if *Y*_*p*_ = *k*_*n*_*/α*_*y*_, *m*_*n*_ always equals to its steady-state value according to Eq. 1).

Based on the simulations in Figure 4b, we noticed that the excess mRNA is degraded exponentially in the reversion phase since *X*_*c*_ has reached its steady value at the beginning of the reversion phase. Therefore, we fit the data during the reversion phase (after 95 minutes) using the following equation:

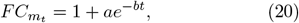

where 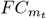 represents the fold change of mRNA concentration relative to the value before perturbation at time *t*, and *a* and *b* are the parameters determined by fitting. On the other hand, since the nuclear mRNA concentration is constant throughout the process, the nuclear mRNA export rate always equals *k*_*n*_. Therefore, we rewrote Eq. 4 as

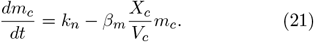

We then integrated Eq. 21 to obtain the temporal changes in cytoplasmic mRNA:

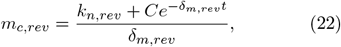

where *m*_*c,rev*_ is the cytoplasmic mRNA number during the reversion phase, *k*_*n,rev*_ is the mRNA production rate during the reversion phase (which is constant in the reversion phase), and *C* is a constant. *δ*_*m,rev*_ is the mRNA degradation rate per mRNA so that 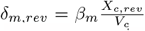, where *X*_*c,rev*_ is the number of *X*_*c*_ during the reversion phase (which is also constant). Given that the amount of nuclear mRNA remains constant, we expressed the fold change of total mRNA as

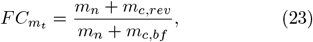

where *m*_*n*_ is the nuclear mRNA number and *m*_*c,bf*_ is the cytoplasmic mRNA number before Xrn1 depletion. Using Eqs. 20-23, it is easy to find that *δ*_*m,rev*_ = *b*.

Experimental data showed that *k*_*n,rev*_ decreased to approximately 0.4 times the mRNA production rate before perturbation *k*_*n,bf*_ : *k*_*n,rev*_*/k*_*n,bf*_ = 0.4 (Figure 4a). Because the cytoplasmic mRNA numbers are the same between the steady states before and after the perturbation, *k*_*n,rev*_*/δ*_*m,rev*_ = *k*_*n,bf*_ */δ*_*m,bf*_. Therefore, *δ*_*m,rev*_*/δ*_*m,bf*_ = 0.4, and *m*_*c,bf*_ = *k*_*n,bf*_ */δ*_*m,bf*_ = 0.4*k*_*n,bf*_ */δ*_*m,rev*_. Typically, the mRNA production rate of budding yeast cells ranges from about 180 to 2300 mRNA per minute [64]. Hence, we set the mRNA production rate of WT cells before perturbation *k*_*n,bf*_ = 500 min^*−*1^, from which we determined the value of *m*_*c,bf*_.

We focused on the accumulation and adaptation phases to estimate the remaining parameters’ values. We rewrote Eq. 13 as:

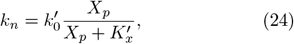

where 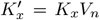, and 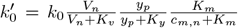. We assumed that 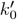 is almost constant throughout the process so that the mRNA production rate *k*_*n*_ (Eq. 13) is approximately proportional to 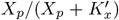 during the depletion experiment of Xrn1.

Calculating the analytical solutions for the time dependence of *X*_*c*_ and *k*_*n*_ was challenging, particularly for *k*_*n*_ as it depends on *X*_*p*_. To simplify the problem, we made the following approximations. Simulations in Figure 4b showed that *X*_*c*_ increases approximately linearly in the accumulation and adaptation phases. Therefore, the transformation speed *v*_*trans*_ = *β*_*x*_*X*_*c*_ from *X*_*c*_ to *X*_*p*_ is also a linear function of time during the accumulation and adaption phase, 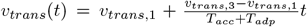. Here, *T*_*acc*_ and *T*_*adp*_ are the durations of the accumulation and adaptation phases. In the steady state before perturbation, *v*_*trans,bf*_ = *k*_*n,bf*_ = *β*_*x*_*X*_*c,bf*_. At the beginning of the accumulation phase, we assumed that 80% of *X*_*c*_ are depleted so that *X*_*c*,1_ *≈* 0.2*X*_*c,bf*_, therefore *v*_*trans*,1_ *≈*0.2*v*_*trans,bf*_. Because the mRNA production rate and the transformation rate must balance as well in the steady after perturbation, *v*_*trans,rev*_ = *k*_*n,rev*_ = *β*_*x*_*X*_*c,rev*_. According to the experimental data, *k*_*n*,3_ = *k*_*n,rev*_ *≈* 0.4*k*_*n,bf*_ at the end of the adaptation phase, so that *v*_*trans*,3_ = *v*_*trans,rev*_ *≈*0.4*v*_*trans,bf*_. Thus, we obtained an approximate expression for the transformation speed. We also calculated the time-averaged transformation rates in the accumulation phase (*⟨v*_*trans,acc*_*⟩*) and the adaptation phase (*⟨v*_*trans,adp*_*⟩*).

Similarly, we approximated the change in *k*_*n*_ in the experimental data with a combination of two linear reductions: a small slope in the accumulation phase and a large slope in the adaptation phase. Likewise, we computed the time-averaged mRNA production rates during these two phases (*⟨k*_*n,acc*_*⟩* and *⟨k*_*n,adp*_*⟩*). With these linear approximations, we calculated the changes in *X*_*p*_ during the accumulation and adaptation phases:

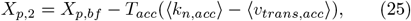

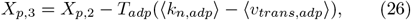

where *X*_*p*,2_ and *X*_*p*,3_ are the numbers of *X*_*p*_ at the end of the accumulation and adaptation phases, respectively. Based on experimental data, the mRNA production rate at the end of the accumulation phase (*k*_*n*,2_) was approximately 0.95*k*_*n,bf*_, while at the end of the adaptation phase, *k*_*n*,3_ was around 0.4*k*_*n,bf*_. Using Eq. 24, we wrote following equations:

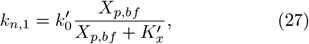

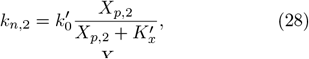

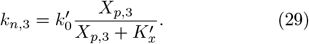

Given that *k*_*n,bf*_ = 500 min^*−*1^, we solved the above equations and obtained the values of 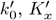, and *X*_*p,bf*_.

Next, we numerically calculated the amount of mRNA accumulated during the accumulation and adaptation phases. Using 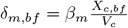, we rewrote Eq. 21 as

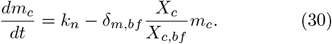

Using the linear approximations of *k*_*n*_ and *X*_*c*_ in the accumulation and adaptation phase, we numerically integrated the equation to obtain the total cytoplasmic mRNA *m*_*c*,3_ at the end of the adaptation phase, where the total mRNA is approximately 1.3*m*_*t,bf*_ (Figure 4a). Therefore,

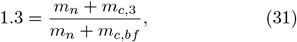

from which we solved for the value of *m*_*n*_.

We have so far obtained the following parameters from the experimental data of [14]: *m*_*c*_ (*m*_*c,bf*_), *m*_*n*_, *X*_*p*_ (*X*_*p,bf*_), 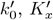, and *δ*_*m*_ (*δ*_*m,bf*_) for a cell without Xrn1 depletion. We next estimated all the parameters as the starting point of the parameter search process. We set *k*_0_ = 6000 *min*^*−*1^, *K*_*y*_ = 100 *f L*^*−*1^, and *Y*_*p*_ = 1*×*10^4^ to solve *K*_*m*_ with the definition of 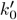. We set *X*_*n*_ = 5000 to solve *α*_*x*_ with Eq. 10. We set *X*_*c*_ = 2 *×* 10^5^ to solve *β*_*x*_ with Eq. 11 and *β*_*m*_ with the definition of *δ*_*m*_. We set *Y*_*p*_ = *Y*_*n*_ = 1000 and solve *α*_*y*_ with Eq. 12. We then solve *α*_*m*_ with Eq. 8. Summing up *X*_*n*_, *X*_*c*_, and *X*_*p*_, we obtained *X*_*t*_ and similarly for *Y*_*t*_.

Finally, we optimized these 11 parameters to fit the experimental data by minimizing the weighted mean squared error (MSE) between the model predictions and the experimental data of the temporal fold changes of mRNA concentration and the mRNA production rates. This optimized set of parameters is the basic set of parameters used in our simulations (Table II).

**TABLE 2.** The basic set of parameters.

| Parameter | Value | Unit |
| --- | --- | --- |
| $k_0$ | 6000.00 | $\text{min}^{-1}$ |
| $K_x$ | 180.18 | $\text{fL}^{-1}$ |
| $K_y$ | 100.00 | $\text{fL}^{-1}$ |
| $K_m$ | 5086.24 | $\text{fL}^{-1}$ |
| $K_v$ | 5.83 | $\text{fL}$ |
| $\alpha_m$ | $3.28 \times 10^{-5}$ | $\text{fL} \cdot \text{min}^{-1}$ |
| $\alpha_x$ | 0.10 | $\text{min}^{-1}$ |
| $\alpha_y$ | 0.52 | $\text{min}^{-1}$ |
| $\beta_m$ | $9.92 \times 10^{-6}$ | $\text{fL} \cdot \text{min}^{-1}$ |
| $\beta_x$ | $2.24 \times 10^{-3}$ | $\text{min}^{-1}$ |
| $X_t$ | $2.30 \times 10^5$ | - |
| $Y_t$ | $1.10 \times 10^4$ | - |
| $V_n$ | 2.80 | $\text{fL}$ |
| $V_c$ | 37.20 | $\text{fL}$ |

### Details of numerical calculations

For the simulations of global genetic perturbations (Figure 2d, e, f, and S4), we calculated the steadystate values of mRNA concentrations, mRNA production rates, and mRNA degradation rates using Eqs. 8-12. We randomly sampled all parameters from lognormal distributions for the blue points in Figure 2d, e, f, and S4. The CVs of parameters involved in the expression of *k*_*n*_, i.e., *k*_0_, *K*_*v*_, *K*_*x*_, *K*_*y*_, *V*_*n*_, and *V*_*c*_, were set to 0.5, while the CVs of other parameters were set to 0.2. The total numbers of X and Y were also randomly sampled from lognormal distributions with a CV of 0.5. For the red points in Figure 2d, e, and S4, we randomly sampled *α*_*m*_ from a lognormal distribution with its CV equal to 1, while the CVs of other parameters in the expression of *k*_*n*_, as well as the total numbers of X and Y, were set to 0.2. In these simulations, we modified *k*_0_ to be 1*/*8 of the value from the set of basic parameters.

For the simulations involving changing cell volumes (Figure 3b, c, and d), we calculated the steady-state values of the mRNA number, the production rate, and the degradation rate per mRNA given a cell volume. We maintained constant concentrations of total X and Y across different volumes. To verify the constant mRNA concentration predicted by the RS model for any volume dependence of the mRNA production rate, we replace the volume-dependence term *V*_*n*_*/*(*<u>V</u>*_*n*_ + *K*_*v*_) in Eq. 13 by 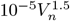 in Figure 3c, and 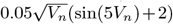 in Figure 3d. In these simulations, we modified *k*_0_ to be 1*/*3 of the value from the set of basic parameters.

For the simulations involving perturbations of Pol II numbers (Figure 3e), we monitored the temporal changes in the mRNA concentration of a specific gene under different values of *K*_*v*_ after silencing the expression of that gene. The parameters used in the simulations (except *K*_*v*_) were optimized from the basic parameters to minimize the squared error (*E*^2^) between the experimental and predicted degradation rates in WT cells (see details in the analysis of experimental data).

For the simulations depicting the temporal changes after acute depletion of Xrn1 (Figure 4a, b, d, e, and S7), we calculated the fold changes of different molecules compared to the respective homeostatic values before depletion. To mimic the acute depletion of Xrn1, we instantaneously removed 80% of *X*_*c*_ from the cytoplasm. In Figure 4d, e, and Figure S7, two simulations are shown together in each plot. All parameters used in these simulations are the same except for the one indicated in each plot.

For the simulations examining temporal changes of mRNA with different concentrations of X or Y (Figure 5a), we monitored the changes of cytoplasmic and nuclear mRNA levels after inhibiting transcription, compared to their values before perturbation. We inhibited transcriptional by instantaneously decreasing the value of *k*_0_ to its 50% at time 0.

For the simulations of temporal changes after complete transcription inhibition (Figure 5c), we introduced a very large *K*_*v*_ to completely shut off transcription at time zero, and then calculated the temporal changes of mRNA and X.

For the simulation of the modified model with additional protein P (Figure 6b), we simulated the temporal changes of the total number of mRNA, the cytoplasmic P (*P*_*c*_ + *P*_*c,b*_), the nuclear P (*P*_*c*_ + *P*_*c,b*_), the mRNA production rate, and the mRNA degradation rate per mRNA using Eqs. 14-18 after increasing the degradation factor *β*_*m*_ by a factor of 10 at time 0, mimicking the globally accelerated degradation of the host mRNA during the viral infection. In these simulations, we modified *α*_*m*_ to be 20 times of the value from the set of basic parameters.

To simulate the extended model involving multiple distinct subsets of mRNAs (Figure 6d), we simulated the transcription of two groups of genes regulated by their own Xs and Ys. The parameters of these two RS models are set the same. We perturbed the expression of one group of genes by decreasing the number of its *X*_*c*_ to its 10% at time zero. We then monitored the temporal changes of the mRNA concentrations of both groups and the total mRNA concentration.

### Finding candidates of X and Y

The RS model suggests that X and Y are multifunctional molecules involved in transcription, exportation, and degradation. To identify candidate genes in these categories, we utilized various databases, including the Gene Ontology (GO) database through AmiGO (version 2.5.17) [65–67], the Kyoto Encyclopedia of Genes and Genomes (KEGG) database [68–70], and the Saccharomyces Genome Database (SGD) [71]. In our search for transcription-related genes, we focused on those annotated as transcription factors or involved in the reg-ulation of transcription, as well as the subunits of Pol II. For exportation-related genes, we looked for annotations related to nuclear transportation factors and the subunits of the nuclear pore. In the context of degradation, we identified genes involved in mRNA catabolism. We marked genes with dual functions in transcription and exportation as Y and genes with dual functions in transcription and degradation as X (Table I).

## Data availability

All data are acquired from public repositories. The experimental data of the global genetic perturbations in mammalian cells were obtained from [17]. The experimental data of the global genetic perturbations in budding yeast cells were obtained from [9]. The experimental data of volume changes were obtained from [26]. The mRNA production rate fold changes were obtained from the Pol II occupancy data in Figure 1e of [26]. Figure 6c of [26] shows the fold change of mRNA degradation rate. The experimental data of perturbing Pol II were obtained from Supplementary Figure 8e of [26]. The experimental data of the acute depletion of Xrn1 were obtained from [14].

## Code Availability

All codes for mathematical simulations are available in the following link (https://github.com/QirunWang/Codes-for-the-RSmodel).

## ACKNOWLEDGMENTS

We thank Yihan Lin for helpful discussions related to this work. All schematics are created with BioRender.com. The research was funded by the National Key R&D Program of China (2021YFF1200500) and supported by grants from Peking-Tsinghua Center for Life Sciences.

