## Supplementary Information for "Homeostasis of mRNA concentrations through coupling transcription, exportation, and degradation"

Qirun Wang[1] and Jie Lin[1,2]
[1] Center for Quantitative Biology, Peking University, Beijing, China
[2] Peking-Tsinghua Center for Life Sciences, Peking University, Beijing, China

### A. Details of the feedback models

In the positive feedback model where mRNA activates its own degradation (Figure S1a), the dynamics of mRNA concentration follows

$$\frac{dc_m}{dt} = k_n - \beta_m c_m^2 \tag{S1}$$

where $k_n$ is the production rate, and $\beta_m c_m^2$ is the degradation rate, a nonlinear mRNA concentration function. We sampled $k_n$ uniformly within a finite range and monitored the relationship between the mRNA production rate $k_n$ and the steady-state mRNA concentration $c_m$ (Figure S1b). We also sampled $\beta_m$ uniformly within a finite range and monitored the relationship between $k_n$ and $c_m$ (Figure S1c). We randomly sampled the mRNA concentrations from a Poisson distribution with the means equal to the steady-state values derived from the positive feedback model, mimicking noises in gene expression. We also added Gaussian noise on top of the mRNA production rates so that the CVs equal 0.003. In neither case we observed mRNA buffering.

In the negative feedback model where mRNA inhibits its own production (Figure S1d), the dynamics of mRNA concentration follows

$$\frac{dc_m}{dt} = \frac{a}{c_m + b} - \beta_m c_m, \tag{S2}$$

where $a$ and $b$ are constants. We sampled $a$ uniformly within a finite range and monitored the relationship between $k_n$ and $c_m$ (Figure S1e). We also sampled $\beta_m$ uniformly within a finite range and monitored the relationship between $k_n$ and $c_m$ (Figure S1f). We randomly sampled the mRNA concentrations from a Poisson distribution with the means equal to the steady-state values derived from the negative feedback model, mimicking noises in gene expression. We also added Gaussian noise on top of the mRNA production rates so that the CVs equal 0.003. In neither case we observed mRNA buffering.

### B. The modified models in which X and Y are not necessary for transcription

We assumed that transcription is accelerated by the binding of $X_p$ to the preinitiation complex (PIC), and transcription can also proceed without $X_p$ (we still kept the necessity of $Y_p$ to transcription for simplicity). We assumed that $X_p$ binds to the PIC and hops off with a binding affinity $K_x$. PICs bound to $X_p$ initiate transcription faster than PICs unbound to $X_p$. Therefore, the mRNA production rate can be decomposed into two parts

$$k_{n,0} = k_1 \frac{K_x}{K_x + X_p/V_n}, \tag{S3}$$

$$k_{n,p} = k_2 \frac{X_p/V_n}{K_x + X_p/V_n}, \tag{S4}$$

where $k_1$ and $k_2$ are coarse-grained variables, including the other factors in Eq. 13 of the maintext. The total mRNA production rate $k_n = k_{n,0} + k_{n,p}$, while the transformation rate form $X_p$ to $X_n$ is $k_{n,p}$. We rewrote Eqs. 1, 4 in the maintext as

$$\frac{dm_n}{dt} = k_{n,0} + k_{n,p} - \alpha_m \frac{Y_n}{V_n} m_n, \tag{S5}$$

$$\frac{dm_c}{dt} = \alpha_m \frac{Y_n}{V_n} m_n - \beta_m \frac{X_c}{V_c} m_c, \tag{S6}$$

(S7)

Meanwhile, $k_{n,p} = \beta_x X_c$ in the steady state to keep a constant number of $X_p$. Therefore, the steady-state solution of the cytoplasmic mRNA concentration becomes

$$c_{m,c} = \frac{\beta_x}{\beta_m} \frac{k_1 K_x V_n + k_2 X_p}{k_2 X_p}. \tag{S8}$$

In this modified model, the cytoplasmic mRNA concentration depends on the cell volume and the copy number of $X_p$: the mRNA buffering breaks down. Numerical simulations confirmed our predictions (Figure S2). A similar analysis also applies to Y. In conclusion, the validity of mRNA buffering requires that proteins X and Y must be necessary factors for transcription initiation.

#### C. The modified model in which Y shuttles

In this modified model, the dynamics of protein X is the same as in the original model, while the dynamics of Y follows

$$\frac{dY_n}{dt} = k_n - \alpha_y Y_n, \tag{S9}$$

$$\frac{dY_c}{dt} = \alpha_y Y_n - \beta_y Y_c, \tag{S10}$$

$$\frac{dY_p}{dt} = \beta_y Y_c - k_n, \tag{S11}$$

where $Y_c$ is the cytoplasmic Y. In this modified model, mRNA buffering is still valid (Figure S3).

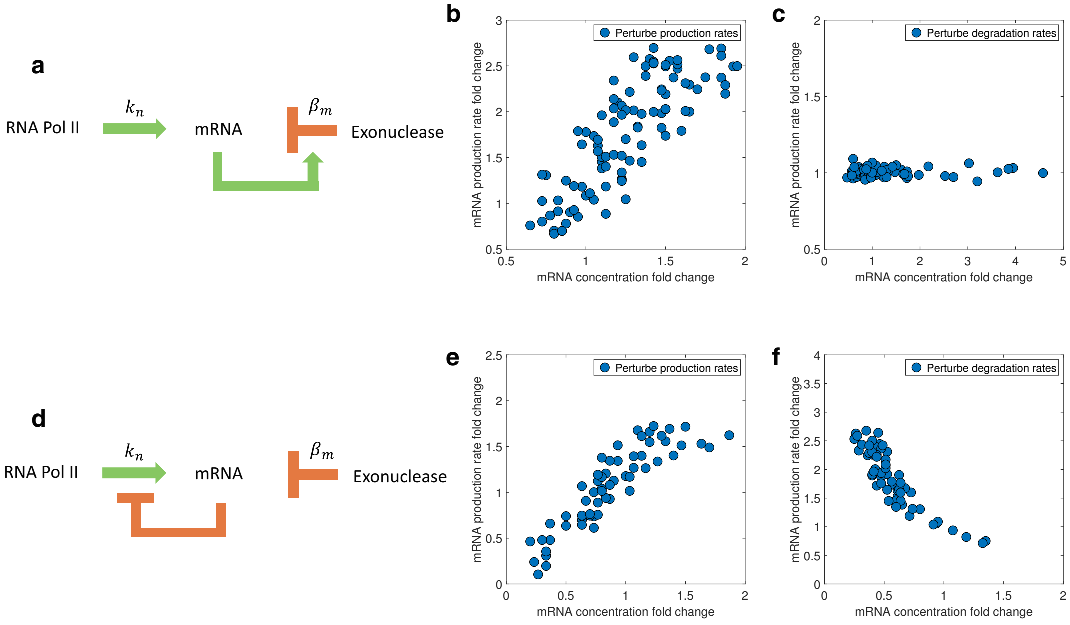


FIG. S1: **Models of mRNA feedback cannot achieve mRNA buffering.** (a) Schematic of the positive feedback model. The green arrow represents activation, and the red blunt arrow represents inhibition. (b) In the positive feedback model, a positive correlation exists between the mRNA production rate and mRNA concentration when the mRNA production rate is perturbed. (c) In the positive feedback model, a zero correlation exists between the mRNA production rate and mRNA concentration when the mRNA degradation rate is perturbed. (d) Schematic of the negative feedback model. (e) In the negative feedback model, a positive correlation exists between the mRNA production rate and mRNA concentration when the mRNA production rate is perturbed. (f) In the negative feedback model, a negative correlation exists between the mRNA production rate and mRNA concentration when the mRNA degradation rate is perturbed. In all scenarios, the mRNA concentration is not buffered.

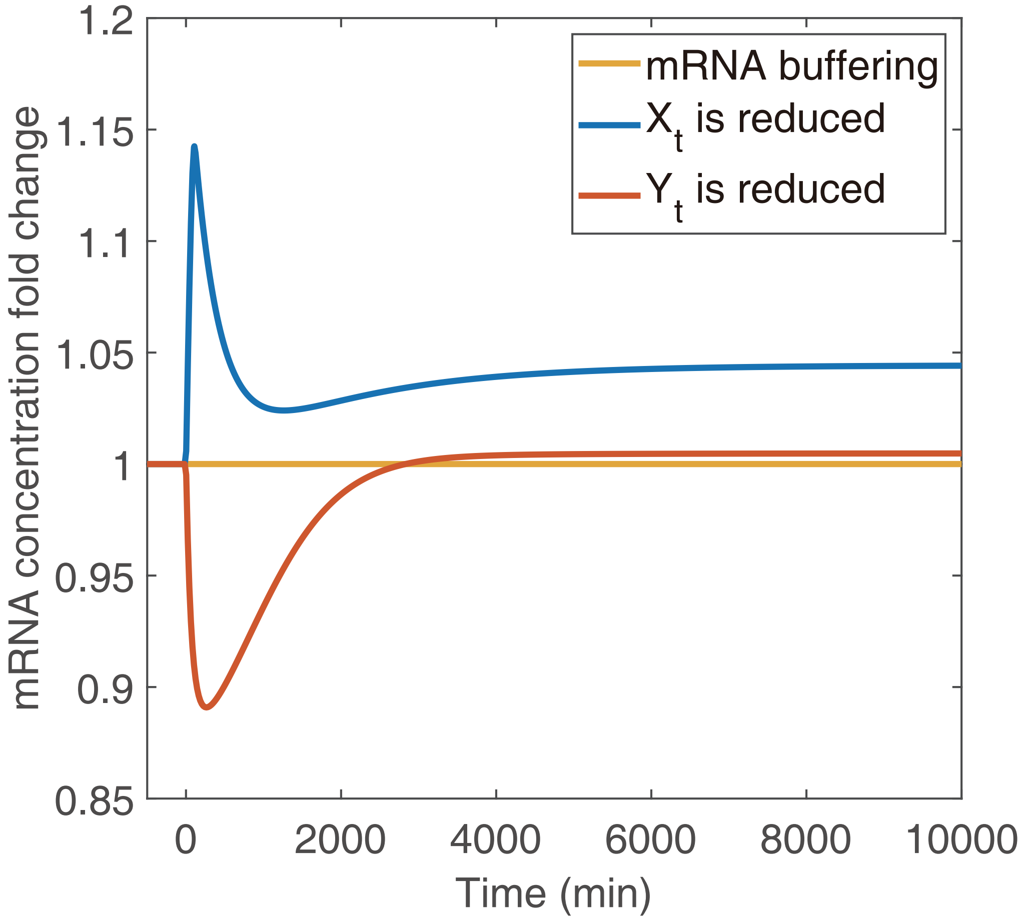


FIG. S2: **Breakdown of mRNA buffering in the modified model where X is not necessary for transcription.** We simulate the modified model in which two kinds of different $k_n$ (Eq. S3 and S4) are introduced. At time 0, we deplete either $95\%X_t$ or $99\%Y_t$, and monitor the temporal changes of the total mRNA levels.

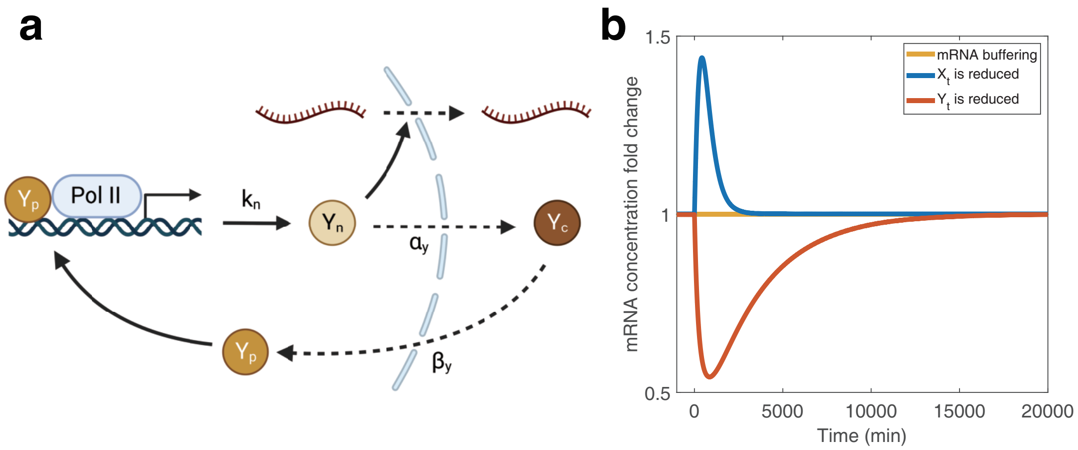


FIG. S3: **mRNA buffering is still valid in the modified model where Y also shuttles.** (a) Schematic of the modified model. (b) The temporal change of mRNA concentration after the total number of X or Y is reduced. We simulated the modified model in which the dynamics of Y follows Eq. S9-S11. At time 0, we depleted either 90%$X_t$ or 90%$Y_t$, and monitored the temporal changes of the total mRNA levels.

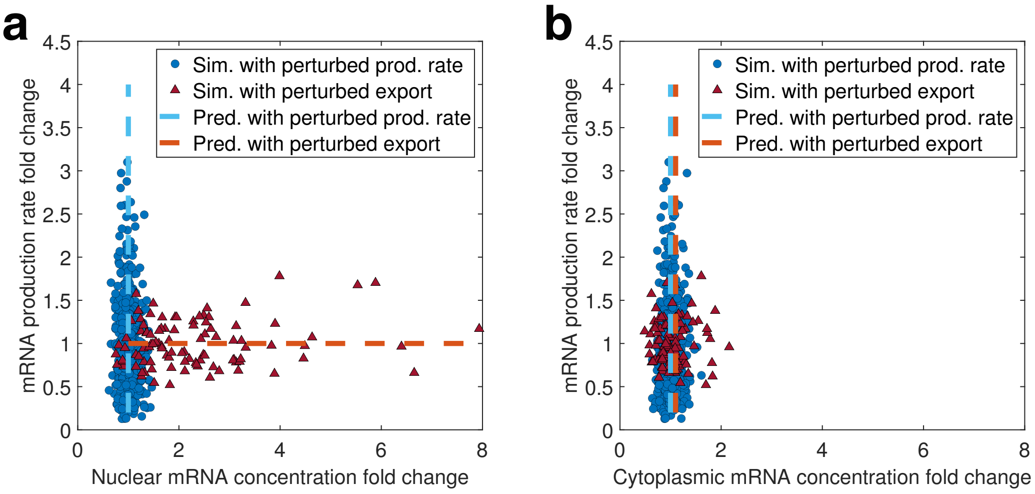


FIG. S4: **Simulations of the relationships between the mRNA production rates and the mRNA concentrations without the negative feedback of nuclear mRNA on transcription.** Sim., simulation; prod., production; pred., prediction. The blue dashed line is an $x = 1$ line, and the red dashed line represents the prediction when $\alpha_m$ is perturbed. (a) Simulations for the nuclear mRNA concentration. The blue circles represent simulation results with multiple parameters perturbed, and the red triangles represent simulation results with $\alpha_m$ perturbed. The same meanings of points apply to (b). We randomly sampled the mRNA copy numbers from a Poisson distribution with the means equal to the predictions of the RS model, mimicking noises in gene expression. We also added Gaussian noise on top of the mRNA production rates so that the CVs equal 0.01. The same noises were also applied to (b). (b) Simulations for the cytoplasmic mRNA concentration.

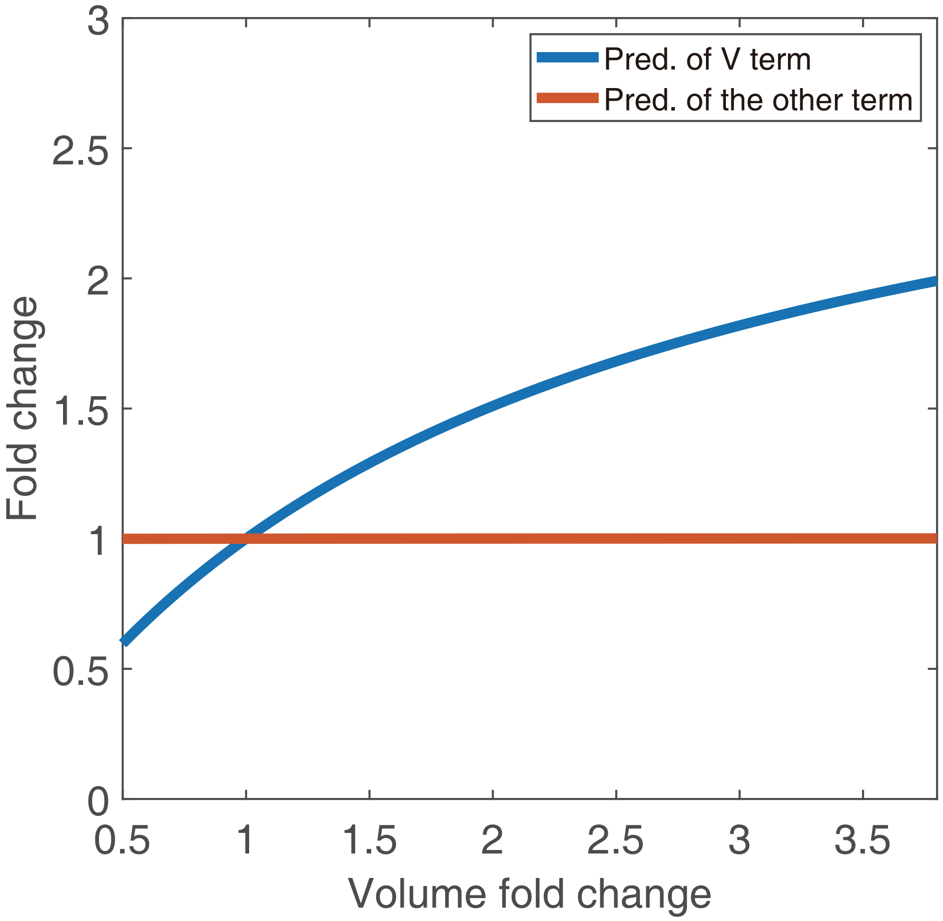


FIG. S5: **The volume dependence of the two terms in $k_n$.** Pred., prediction. Here, the V-term is $\frac{V_n}{V_n+K_v}$, and the other term is $k_0\frac{x_p}{x_p+K_x}\frac{y_p}{y_p+K_y}\frac{K_m}{c_{m,n}+K_m}$. The lines are from the same simulations as Figure 3b in the maintext.

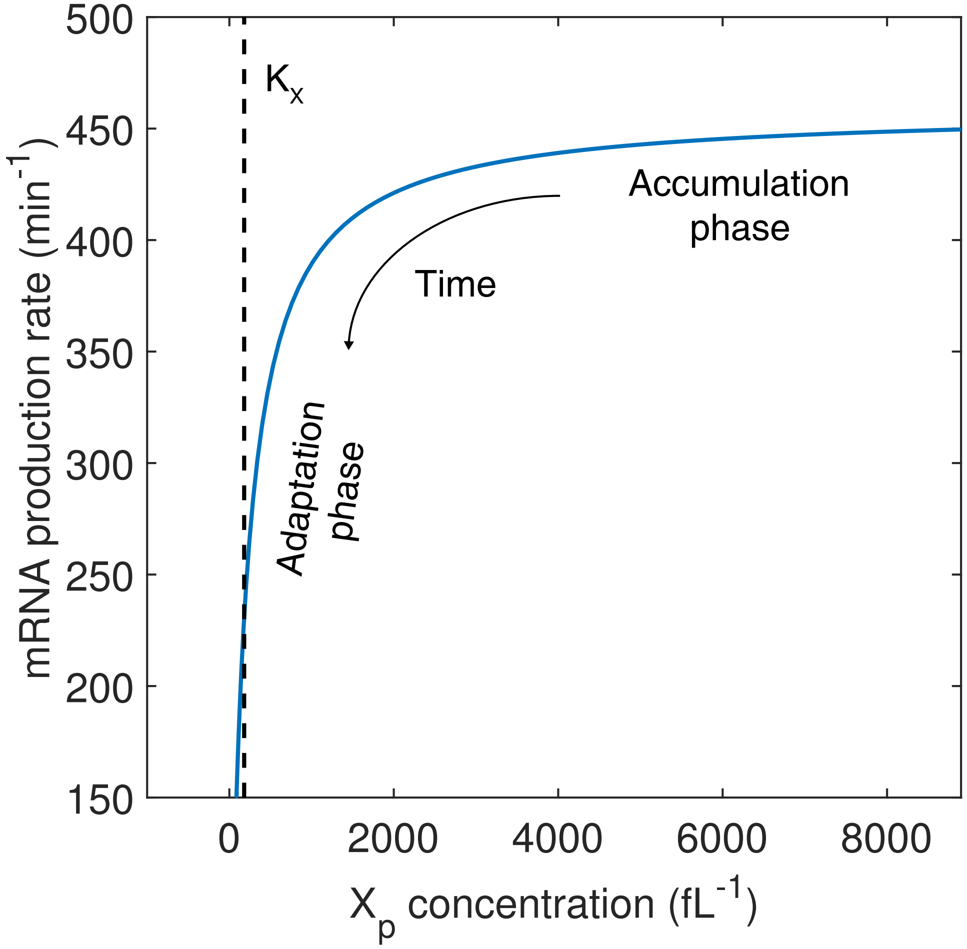


FIG. S6: **Relationship between $X_p$ and mRNA production rate in the simulation.** $X_p$ initially decreases with a nearly constant rate until it hits the Michaelis-Menten constant $K_x$, which triggers a significant decrease in the mRNA production rate.

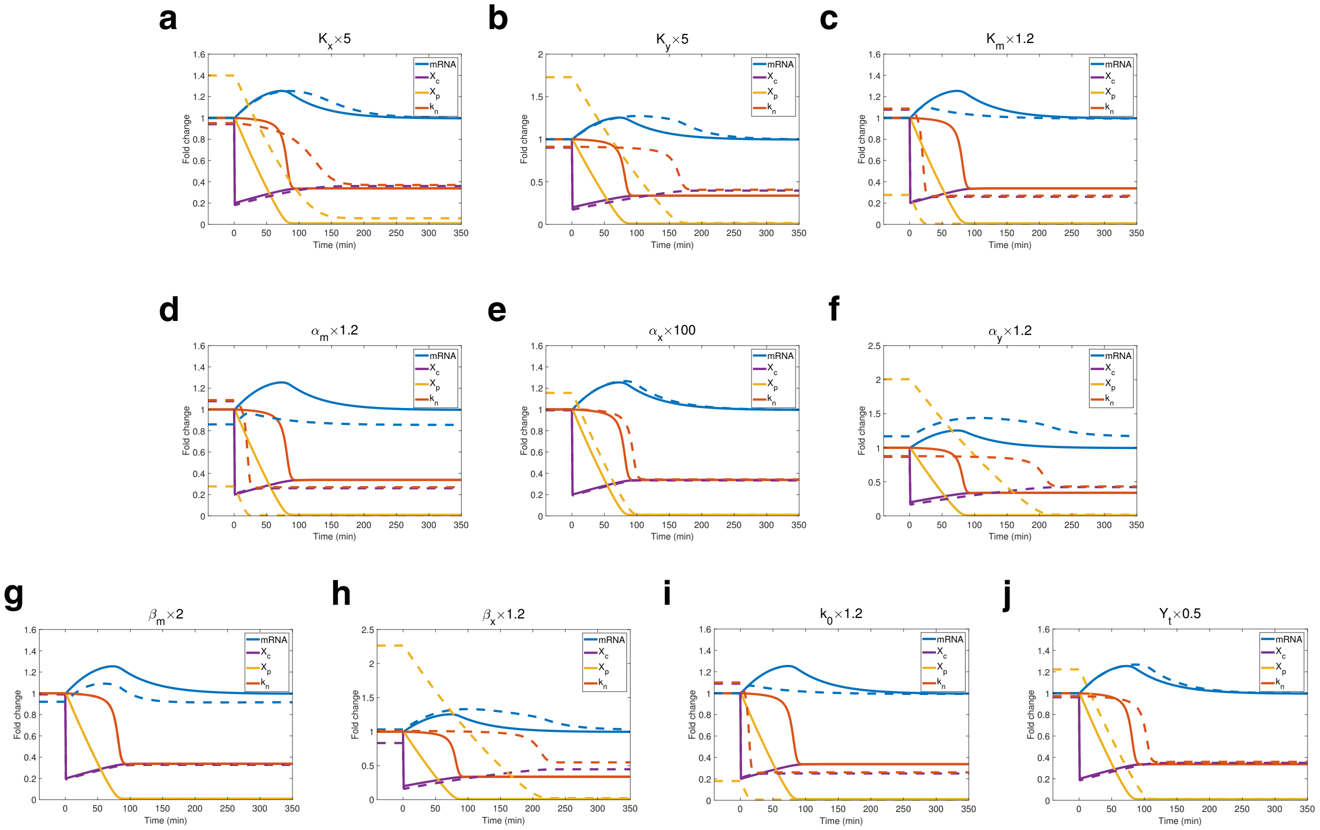


FIG. S7: **Influence of different parameters on the timescales of temporal changes after depleting $X_c$.** The titles show how parameters are manipulated compared to those in WT cells. Solid lines show the temporal changes of WT cells, while dash lines show the temporal changes of cells in which parameters are manipulated. Both the solid lines and the dashed lines are the fold changes relative to the homeostatic values of the WT cells before depletion.

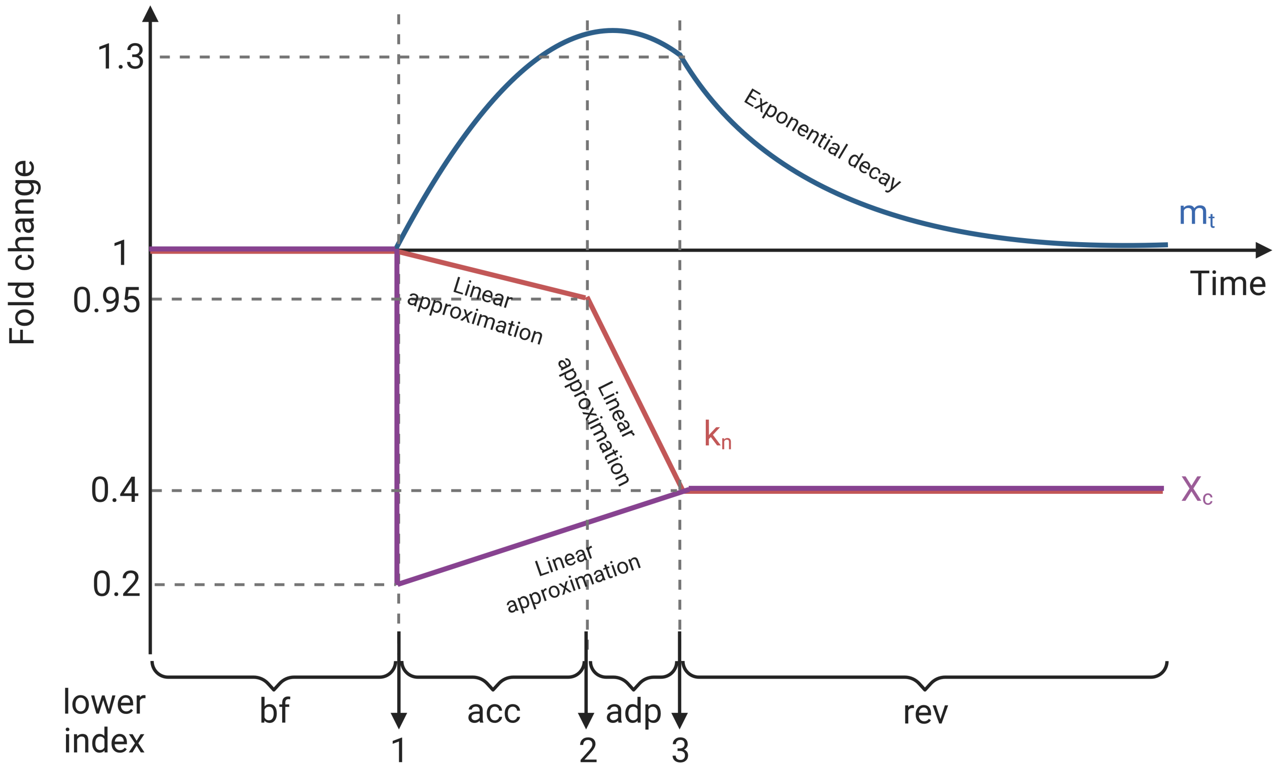


FIG. S8: **Simplification of the transcription dynamics to determine fitting parameters.** We used an exponential decay model to fit parameters during the reversion phase. We used linear approximations to model the dynamics of $X_c$ and $k_n$ during the accumulation and adaptation phases. The lower indexes used in Methods at different times are shown at the bottom of the schematic.
